# Cellular timing heterogeneity regulates phase transitions in living matter

**DOI:** 10.64898/2026.06.16.732542

**Authors:** Magdalena Schindler-Johnson, Jef Vangheel, Adrián Aguirre-Tamaral, Tom E. R. Belpaire, Bart Smeets, Bernat Corominas-Murtra, Nicoletta I. Petridou

**Author notes:** These authors contributed equally (Magdalena Schindler-Johnson, Jef Vangheel).

## Abstract

Rigidity transitions govern tissue organization in ways reminiscent of inert materials. Yet, living tissues are composed of active units with autonomous timing mechanisms, raising the question whether microscopic cellular timing distribution influences collective mechanical states. Here we identify heterogeneity in cellular timescales as a heritable parameter, regulating rigidity transitions in embryonic tissues. Lineage tracking and quantitative mechanical analysis reveal that zebrafish morphogenesis starts with a tissue rigidity collapse occurring at maximal cell cycle length heterogeneity. This heterogeneity arises from size-dependent stochastic differences in resource allocation, with resource availability defining the cell cycle length. Such differences are inherited across generations, amplifying and structuring tissue-wide cell cycle length heterogeneity. Experiments and large-scale 3D simulations identify an optimum level of cellular timing variability at which cell-cell contact remodelling is spatially coordinated driving timely and robustly the rigidity transition. These findings demonstrate that embryos exploit microscopic temporal disorder for timing and tuning tissue morphogenesis.

---

Biological tissues are increasingly understood as living materials, whose deformation and organization can be predicted from local interaction rules ^1-3^. A growing body of work at the interface of developmental biology and soft active matter has traced these rules to simple cellular properties –like density, connectivity, adhesion and tension fluctuations– acting as control parameters of tissue material phase transitions, and driving spatiotemporal shifts in rheological properties during development and disease ^4-8^. However, living tissues challenge key assumptions underlying the behaviour of inanimate matter: Cells are active agents that continuously consume energy ^9^, are intrinsically heterogeneous ^10-15^, and, crucially, operate on autonomous timescales due to internal biochemical clocks ^16^, metabolic rates ^17^ and biological inheritance ^18-20^. Whether such cellular timing distributions shape the emergent rheology and organization of living tissues remains largely unresolved.

Developmental programs regulating cellular timing, like cell proliferation, are fundamental drivers of tissue material changes ^21,22^. Cell division can promote both tissue fluidization or solidification, either via inducing contact remodelling ^23-27^ or crowding ^28-30^, respectively. During tissue fluidization, different ways of division-dependent contact remodelling have been suggested based on tissue confluency. In non-confluent tissues, mitotic rounding during the cell cycle reduces cell-cell contact size and overall cell connectivity ^6,23^, whereas in confluent tissues, cell divisions induce T1 transitions and topological changes, allowing neighbour exchanges ^24-27^. Division events are heterogeneous in space and time, and heterogeneity in division timing is key for systems-level phenomena, including symmetry breaking, patterning and morphogenesis ^12,31-34^. How cellular timing heterogeneity shapes the form and function of active matter remains unclear, in part because most theoretical approaches –largely developed for non-proliferative systems– treat biological stochasticity as analogous to thermal noise which is frequently externally imposed ^35-38^. However, in proliferative active matter ^39^, duplication rates depend on energy consumption ^40^, are regulated by intrinsic biochemical clocks ^41^, and cell cycle length and phase durations are transmitted along lineages ^19,20,42,43^. This suggests that variability in division timing may follow fundamentally different, lineage-structured dynamics, with potentially distinct consequences for collective mechanical behaviour. Experimentally, this problem is further complicated by the difficulty of tuning cellular timing variability independently of mean or average cell dynamics, limiting direct access to the role of cellular temporal disorder in tissue organization. Here, we integrate stochastic modelling, targeted experiments, and large-scale 3D simulations to derive the origin and dynamics of cellular timing variability from basic biochemical and biophysical principles, and to mechanistically decouple it from mean behaviour. We use as a paradigm the cell cleavage division timing, which is a primary source of cellular heterogeneity in early embryogenesis ^22^, and examine how it affects tissue mechanical transitions in zebrafish. We identify an optimum level in division time variability driving a bidirectional tissue material phase transition, enabling embryo morphogenesis. This optimum emerges from stochastic differences in resource allocation amplified by deterministic hyperbolic growth of the cell cycle length ^44^. The amplification generates lineage-inherited variability, resembling a memory effect, which controls the temporal and spatial pattern of cell-cell contact remodelling, and underlies the robustness and stereotypicality of the tissue mechanical transition at the onset of embryo morphogenesis. Our findings uncover structured, heritable temporal noise as a regulator of active matter phase transitions in embryonic tissues.

## Cell cycle length variability peaks at a tissue rigidity collapse

To investigate the interplay of cellular timing heterogeneity and tissue material phase transitions we used a well-established *in vivo* system, the zebrafish early embryo, which initiates its morphogenesis with a cell division-driven tissue fluidization ^23^ (**Fig. 1A**). Zebrafish blastoderm fluidization was shown to be stereotypical, occurring exactly at the onset of its spreading over a yolk cell, followed by tissue rigidification by gastrulation stage ^6,23^. Previous studies showed that this fluidization depends on cell divisions, specifically, on cell-cell contact remodelling occurring during mitotic rounding (**Fig. 1B**). Intriguingly, cell divisions are highly heterogeneous during this time due to the phenomenon of cell cleavage desynchronization ^45^. Across metazoans, upon fertilization, cell cycles are rapid and synchronous until the so-called Mid-Blastula Transition (MBT), where cell cycles lengthen and start varying between the cells, becoming asynchronous by the onset of morphogenesis ^46,47^ (**Movie 1, Fig. 1C**). In consequence, to explore how cell cycle length (CCL) heterogeneity is implicated in tissue material properties, we first characterized the joint spatiotemporal dynamics of CCL dynamics and tissue rigidity during the blastula to gastrula stages in zebrafish embryos.

**Figure 1.**
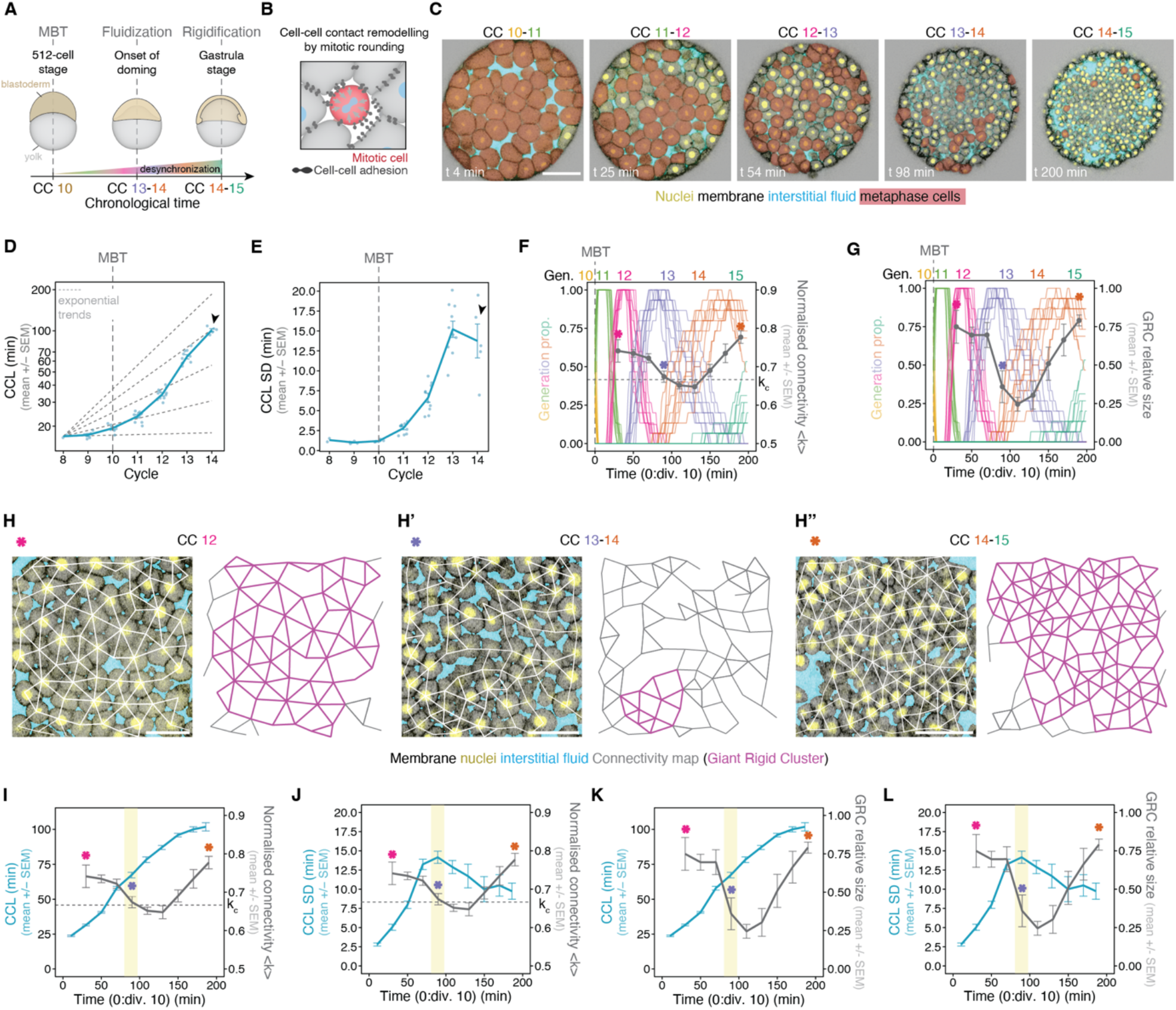
A tissue rigidity transition occurs at a peak in cell cycle length variability. **(A)** Scheme depicting zebrafish development from MBT to gastrula-stage indicating the cell cycle (CC) number counting from fertilization. **(B)** Schematic depiction of the effect of mitotic rounding on cell-cell junctions. **(C)** Example images of a transgenic embryo expressing histone (actb2:H2B-GFP) and membrane markers (actb2:Lyn-tdTomato) injected with dextran-647 in the interstitial space, and imaged from MBT (CC 10) onwards. Red shaded areas indicate metaphase cells. **(D, E)** Mean (D) and standard deviation (SD) (E) of cell cycle length (CCL) per embryo (n=740 cells, N=10 embryos). Black arrowheads indicate plateau of CCL (D) and drop of CCL SD (E), respectively. Dashed lines show different exponential trends for CCL from CC 8 onwards **(F, G)** Overlay of (left y-axis) generation plot showing the proportion of cells at the indicated generation per embryo over time (each line represents one embryo, n=940 cells, N=10 embryos) with (right y-axes) average network connectivity (F) and Giant Rigid Cluster (GRC) size (G) (n=48 networks, N=3 embryos) binned by 20 min intervals. Horizontal dashed grey lines indicate the connectivity critical point k_c_. **(H-H’’)** Exemplary confocal sections of the central blastoderm (left), overlaid with their connectivity networks and marked GRC in purple (right), at the indicated stages. **(I, J)** Plots of overlaid CCL mean (I) and CCL SD (J) (n=666 cells, N=10 embryos) with network connectivity (same data as in F). Horizontal dashed grey lines indicate the connectivity critical point k_c_. (**K, L**) Plots of overlaid CCL mean (K) and CCL SD (L) (same data as in G, J) with GRC size (same data as in G). Asterisks in (H-H’’) indicate the developmental time at which networks were analysed from (E) and (G) (magenta, generation 11-12; purple, generation 13; orange, generation 14-15). The yellow shaded region in (I-L) indicates the peak in CCL SD which coincides with tissue fluidization. (D-G and I-L) error bars indicate mean +/-SEM. CC, Cell Cycle; CCL, Cell Cycle Length; Div., Division; GRC, Giant Rigid Cluster; MBT, Mid-Blastula Transition; SEM, Standard Error of the Mean; SD, Standard Deviation. Scale bars: (C) 100 µm, (H-H’’) 50 µm.

We performed live imaging of embryos labelled for membrane, histone and interstitial fluid (intercellular gaps) across cleavage cycles spanning the synchronous to asynchronous periods ^48^ (**Fig. 1C**). We staged embryos by cell size, since divisions are reductive and daughter cells have roughly half the size of the mother cell (**Fig. S1A**), and quantified CCLs as the metaphase-to-metaphase time difference. In agreement with previous reports ^42,45,49^, we observe that cell cleavages are relatively fast (∼17min) until generation 10 (**Fig. 1D**), which corresponds to the pre-MBT period. Upon MBT, the cell cycle sharply elongates in a faster than exponential manner (**Fig. 1D**). Quantification of the cell-to-cell variability in CCLs, expressed both as the standard deviation (SD) or the coefficient of variation (SD/mean), confirmed that, before MBT, cell cycles are synchronous and variability is low whereas, after MBT, while the cell cycles elongate and desynchronize, variability is rapidly increasing (**Fig. 1E, Fig. S1B**). Moreover, the tissue displays a peak in CCL variability at the 13^th^ generation, which drops right afterwards (**Fig. 1E, Fig. S1B**, arrowheads), since during this time the CCL starts to plateau (**Fig. 1D**, arrowhead). As a result, although cells start exhibiting more similar CCLs after the 13^th^ generation, they appear totally asynchronous when observed under chronological time, resulting in generational overlap (**Fig. S1C, D**). The timing of the peak of CCL variability is close to the onset of blastoderm spreading when tissue fluidization occurs, at around generation 13 ^23^ (**Fig. 1A**, onset of doming).

To simultaneously monitor changes in cell division timing and in the tissue mechanical state, we used rigidity percolation theory, a theoretical framework that predicts the tissue rheological response from parameters extracted from imaging data, specifically cell-cell connectivity ^6,8^. We abstract 3D embryonic tissues as 2D cellular networks, in which the cell centroid is a node and the cell-cell contacts are edges with viscoelastic properties ^2,6^ (**Fig. 1H-H’’**). Within this framework, the material response of the network can be quantified by the size of the largest connected cluster within which nodes have no independent movements, the Giant Rigid Cluster (GRC) (**Fig. 1H-H’’**, purple cluster) ^50,51^. The size of the GRC grows abruptly, from a negligible fraction of the network to spanning almost the entire tissue, when the average connectivity <*k*> crosses a critical point, the isostatic point for Maxwellian rigidity ^6,52^ (**Fig. S1E, E’**), corresponding to ∼4 contacts per cell (<*k*_*c*_> ≈ 0.67, normalized to the maximum achievable connectivity). This change in the topological properties of the tissue induces, in turn, a sudden change in its rheological properties, i.e., fluid-like (subcritical <*k*>) vs. solid-like (supercritical <*k*>) ^2,6^. Applying this network analysis to the same embryonic tissues used to quantify cell cycle dynamics, revealed that the initial crossing of the connectivity critical point, and the corresponding collapse of the GRC, coincides with the 13^th^ generation (**Fig. 1F, G**, purple asterisks) and aligns precisely with the peak of CCL variability (**Fig. 1I-L, Fig. S1F, G**, yellow bands).

The temporal concurrence of the above events, together with previous work showing that cell division is a developmental regulator of tissue mechanical transitions ^23,24^, raises the question whether the timing and nature of the tissue rigidity transition are not only determined by the cell duplication rate, but also its heterogeneity. This raises the hypothesis that the population-level cell cycle heterogeneity could act as a mechanical timer for tissue phase transitions in embryogenesis.

### Hyperbolic resource-dependent division dynamics lead to amplification and inheritance of cell cycle variability

In order to address if and how cell cycle heterogeneity regulates the mechanical transition, we sought, first, to uncouple cell cycle lengthening from CCL variability. To do this, we first explored the underlying mechanisms of cell cycle lengthening and its variation. Cleavages are reductive divisions, composed of S and M phases, which are synchronous before MBT, exhibiting spatial patterns of fast mitotic waves ^53-55^. At MBT, the cell cycle elongates, and G1/G2 phases are incorporated ^56^. Several molecular mechanisms have been proposed to trigger cell cycle elongation, suggesting that cell-to-cell differences in such mechanisms might be responsible for the observed variability. These include titration of maternally deposited materials (e.g., replication factors, histones, dNTPs) ^57-62^, exhaustion of energy reservoirs (e.g., metabolites, lipid droplets) ^63,64^, changes in translational and post-translational activity of the cell cycle regulators ^65-67^, zygotic genome activation ^68^ and many others ^69^. Although the above mechanisms are not conserved across species, there is a consensus that at a given nucleus-to-cytoplasmic ratio (N/C), reflecting the concentration of maternal resources relative to the nucleus content, cells start to elongate their cycles in a cell size-dependent manner, with smaller cells exhibiting longer cell cycles (**Fig. S2A**) ^45,55,70-76^. As a result, differential resource allocation between daughter cells could be responsible for the observed variability. To test this hypothesis, we generated embryos that lack cell boundaries on one side, while the other side retains intact cell compartments. In the former case, the tissue becomes a syncytium where nuclei share resources, whereas in the latter case, each cell divides based on the amount of resources that is enclosed in its compartment, which is ultimately determined by its cell size. To generate this partial syncytium, we injected an Aurora B kinase inhibitor to block cytokinesis within a small cell population ^77^ (**Fig. S2B, C, Movie 2**). Quantifying CCL dynamics post-MBT revealed that the average CCL was unchanged in the partial syncytium, however, its CCL variability did not increase and stayed at comparable levels as pre-MBT cycles (**Fig. S2D, E**). These findings suggest that differential resource allocation between daughter cells accounts for CCL variability.

To understand how resource allocation defines CCL and its variation, we closely quantified their dynamics. Concerning CCL elongation, it appears to grow at a faster than exponential pace (**Fig. 1D**), resembling a hyperbola. Hyperbolic growth is characterized by a rapidly accelerating growth towards a mathematical singularity (a finite point where the value becomes infinite) ^78^ (**Fig. 2A**). This implies a decay of the duplication speed (inverse of CCL), where the singularity would correspond to the “touching” of the zero value, meaning infinite CCL (**Fig. 2B**, red asterisk). Indeed, the duplication speed linearly decays from MBT onwards and would reach zero at the 14^th^ generation if the cell cycles were to elongate with the same dynamics (**Fig. 2B**, dashed line leading to red asterisk). Moreover, in contrast to the pre-MBT cell cycles, where CCL variability increases slowly, resembling a stochastic drift ^12,55^, post-MBT, CCL variability rises rapidly (**Fig. 1E**). However, given that the embryo approaches a maximum CCL at the 14^th^ generation, both CCL and the corresponding variability drop (**Fig. 1D, E, 2B, E, F** arrowheads). In fact, the above CCL variability dynamics deviate from those expected for a typical Brownian drift ^79^, which follow the 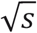 growth of the SD, being *s* the generation (**Fig. S2F**). This suggests that CCL variability is not just the outcome of the accumulation of stochastic differences in resource allocation. Instead, it points to the existence of noise amplification dynamics, increasing variability much more than the one expected by a standard Brownian drift.

**Figure 2.**
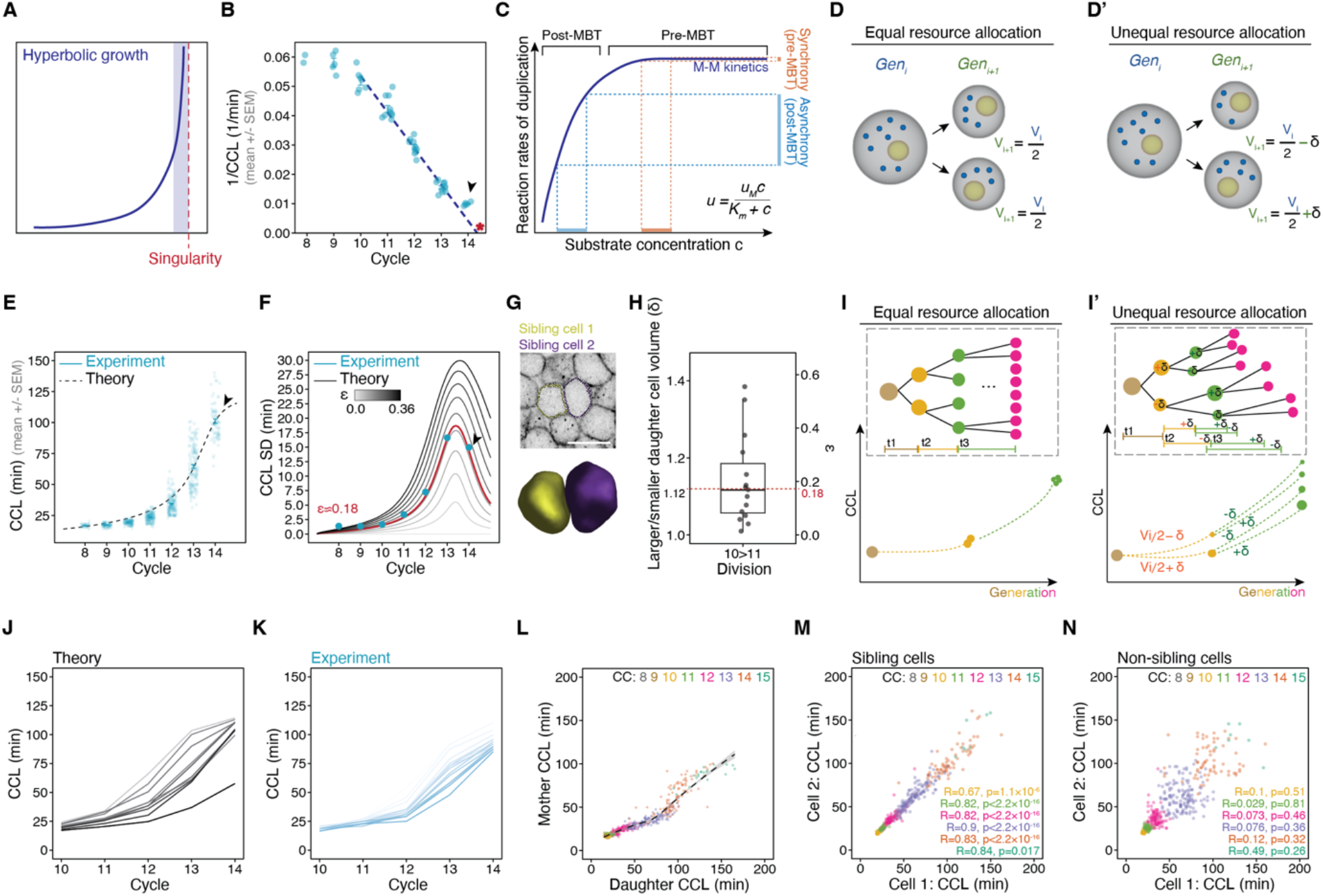
Cell cycle length variability arises by coupling deterministic hyperbolic cell cycle elongation to stochastic resource allocation. **(A)** Schematic depiction of hyperbolic growth behaviour highlighting the singularity that is asymptotically approached at a finite time. The opaque blue band indicates the interval where the function grows very quickly towards infinity as the x-axis approaches the singularity. **(B)** Cell division rate per embryo (1/CCL) embryo (n=740 cells, N=10 embryos) overlaid with theoretical prediction of CCL mean (dashed line). Singularity is marked by the red asterisk. **(C)** Schematic depiction of the Michaelis-Menten-like (M-M) relationship of cell duplication rate *u* and substrate availability *c* during reductive cleavages. Pre-MBT, high substrate availability allows maximal duplication rates *u*_*M*_, resulting in synchronous cell divisions. Post-MBT the substrate is limited, leading to a linear relationship between *c* and *u*, resulting in asynchronous divisions. The formula shows M-M kinetics, where *u*_*M*_ represents the maximum reaction rate and *K*_*m*_ a kinetic constant. **(D, D’)** Scheme depicting the effect of equal (D) and unequal resource allocation (D’) due to cell volume differences δ. **(E-F)** Pooled CCL mean and SD (n=740 cells, N=10 embryos) overlaid with theoretical predictions of the CCL mean (E), and CCL SD at different values of noise *ε* (F). Arrowhead indicates plateau of CCL. **(G)** Exemplary confocal section (top) and 3D volume (bottom) showing segmentation of a sibling cell pair following the 10^th^ division. **(H)** Ratio of volumes of larger to smaller daughter cell (n=15 cell pairs, N=4 embryos; dashed line indicates median volume ratio of 1.116, corresponding to *ε* = 0.18). **(I)** Scheme depicting the effect of equal (I) or unequal (I’) resource allocation on cell division timing and CCL distribution throughout consecutive cell divisions. (**J**) Theoretical prediction of CCL in different cell lineages at indicated CC (n=20 lineages). Colour indicates condition and intensity indicates cell lineage. (**K**) CCL in different lineages within one embryo, separated since at least division 9 over indicated cleavage rounds (n=15 lineages, more replicates in Fig. S3G, G’). (**L**) Correlation of CCL in mother and daughter cells at indicated CC (n=622 cells, N=10 embryos; dashed line is LOESS fit +/-SE, Pearson correlation: R=0.96, p<2.2 × 10^−16^). (**M-N**) Pairwise comparison of CCL of sibling (M) and non-sibling cells (N) (n=1393 cells, N=3 embryos). Statistics: two-sided Pearson correlation. CC, Cell Cycle; CCL, Cell Cycle Length; Gen, Generation; MBT, Mid-Blastula Transition; V, Volume; SD, Standard Deviation; SE, Standard Error. Scale bar: (G) 50 µm.

To elucidate how the hyperbolic dynamics arises, we derived a mathematical model based on the assumptions that (i) maternal resources needed for duplication are being depleted, and (ii) the reaction rates of resources acting as substrates for cell duplication follow fundamental principles of chemical reactions, specifically Michaelis-Menten (M-M) dynamics ^80,81^ (validated in ^44^). In particular, M-M dynamics states that the speed of the chemical reactions *u* non-linearly depends on finite resource/substrate concentration *c* in such a way that reaction rates are fast and approximately constant in the presence of enough resources, but slow down dramatically when the substrate concentration decreases ^80^ (**Fig. 2C**). Specifically, the reaction rate *u* depends on the concentration as:

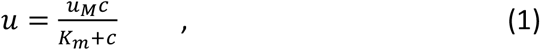

being *u*_*M*_ and *K*_*m*_ the maximum reaction speed and the M-M constant, respectively. Maternal resources are enclosed within the first cell, the zygote, and in each cleavage, although the nuclei content stays the same, the cytoplasmic content is being consumed, since there is no or little synthesis of new resources at these stages ^44,82,83^ (**Fig. 2D**). This implies that, as cells become smaller, the net amount of resources decreases further, leading to an increasingly faster concentration drop. Based on this, we mathematically derive the evolution of the CCL throughout generations, leading to a hyperbolic expression for the CCL growth, with a clear match to the experimental data ^44^ (**Fig. 2E**, see Supplementary Theory Note). These results imply that cell duplication is defined across a biochemical timescale, dictated by the availability of finite maternal resources that act as a substrate for the cell cycle, which, under the M-M dynamics, the duplication rate is expected to slow down dramatically (**Fig. 2C**). This effectively maps pre-MBT cycles to larger cells and resource abundance and post-MBT cycles to smaller cells and resource restriction (**Fig. 2C**).

Building on the above model, we further hypothesized that CCL variability may emerge from intrinsic biological noise that gets amplified by the underlying hyperbolic dynamics. Specifically, unavoidable stochastic cell-to-cell volume differences due to not exactly precise division into two equal daughter cells would lead to heterogeneous resource allocation (**Fig. 2D’**). We mathematically implemented this by introducing a small stochastic element in the volume of daughter cells under the assumption that cell volume approximates available resources (**Fig. 2D’**). This leads to the following stochastic differential equation for the dynamics of the CCL elongation *L* as a function of the generation *s* (see Supplementary Theory Note for derivation details):

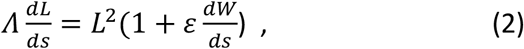

where *Λ* is a constant arising from the M-M constant and the maximum reaction rate, *ε* the noise amplitude and *dW* the standard Brownian differential. This equation leads to a singular behavior and therefore, it is integrated under the constraint that there is a maximum CCL, imposed via a sigmoid-like multiplying factor whose threshold value is extracted from the experimental data (**Fig. 1D**, arrowhead, see Supplementary Theory Note). The best fit to the CCL variability observed in the embryos corresponds to a noise level *ε* ≈ 0.18 in the stochastic dynamical equation (2) (**Fig. 2F**, red curve) (see Supplementary Theory Note). Importantly this noise level is obtained when introducing stochastic variability of around ∼12% of the 3D volume which closely matches the cell size differences measured in experimental data (**Fig. 2G, H**). Overall, this suggests that when the system is under resource saturation (pre-MBT), small stochastic cell-to-cell differences in resource availability do not cause any detectable differences in the duplication rate, thus divisions are synchronous (**Fig. 2C**, orange lines). However, the same differences in resource availability in the regime of low resource concentration (post-MBT) will dramatically amplify the variability in the duplication rate, thus divisions become asynchronous (**Fig. 2C**, blue lines).

Importantly, given that post-MBT cell-to-cell differences in duplication rates are so big, they cannot be resolved through stochastic fluctuations in the next cell cycle rounds. This implies that differences in CCLs will be inherited throughout generations, meaning that the fastest mother cell will give rise to the fastest daughter cells and the fastest granddaughter cells and so on, effectively creating well-separated cell lineages (**Fig. 2I, I’**). We explored whether this behaviour could be explained just by the dynamics of equation (2). At the mathematical level, we generated sibling cells that share the same trajectory until the division event and from then on, their trajectories evolve independently. Analysis of these trajectories showed a clear conservation of ranking regarding the CCL throughout generations, matching those observed in the embryo (**Fig. 2J, K**). The inheritance of CCLs within the lineage is further supported by the strong correlations between the CCL of the mother and daughter cells (**Fig. 2L**) ^42^ and sibling cells (**Fig. 2M**), but not in non-sibling cells (**Fig. 2N**).

Overall, experimental hallmarks of early embryo development including: (i) onset of MBT (**Fig. 2C**), (ii) CCL elongation post-MBT (**Fig. 2E**), (iii) desynchronization post-MBT (**Fig. 2F**), and (iv) correlations between cell size and CCL (**Fig. S2A**) and between cell lineage and CCL (**Fig. 2L-N**) ^20,42,74,75^ are accurately predicted by the above mathematical model. By attributing all these hallmarks to the inheritance of coupled deterministic CCL information and stochastic cell-to-cell differences in resource allocation, these results mechanistically explain the dynamics of CCL elongation and variability of early embryos.

### Inherited cell size heterogeneity defines *in vivo* CCL variability

The above proposed mathematical framework identifies cell volume and cell-to-cell volume differences as the main drivers of CCL evolution and heterogeneity. Since reductive cleavages lead to exponential volume reduction (**Fig. S1A**), stochastic volume fluctuations remain unresolved between smaller and larger cells throughout the generations. Cell size, therefore, becomes an inherited trait throughout the lineage (**Fig. S2G**). To address if in the embryo the natural cell-to-cell size differences are sufficient to account for CCL variability, we designed experiments to manipulate relative cell size differences and examined whether there are effects in CCL variability (**Fig. 3A**). Cell size differences in zebrafish were reported already before MBT due to not entirely symmetric cell divisions ^55,84^. For instance, molecular asymmetries were observed at the 16-cell stage, where the spindle mitotic centrosomes towards the embryo midline are larger in a polo-like kinase (PLK) 1/4-dependent manner ^84^. This potentially leads to slightly asymmetric divisions and may result in larger cells towards the embryo centre (**Fig. 3B**). Indeed, by inhibiting PLK1/4, we were able to reduce cell size differences without affecting the average cell volume (**Fig. 3C, F, Fig. S3A**). The small reduction in the variability of cell sizes was sufficient to reduce CCL variability, without affecting the mean CCL (**Fig. 3G, Fig. S3D**). Conversely, to increase relative cell size differences, we performed micromanipulations in one-cell stage embryos to increase cell size differences between the first two sibling cells. Using a glass micropipette, we removed material containing cytoplasmic and yolk components ^85^, either symmetrically, at the centre of the embryo, or asymmetrically, at one side of the embryo (**Fig. 3D**). The latter resulted in an asymmetric first cell cleavage, generating daughter cells of different sizes (**Fig. 3D-F**).

**Figure 3.**
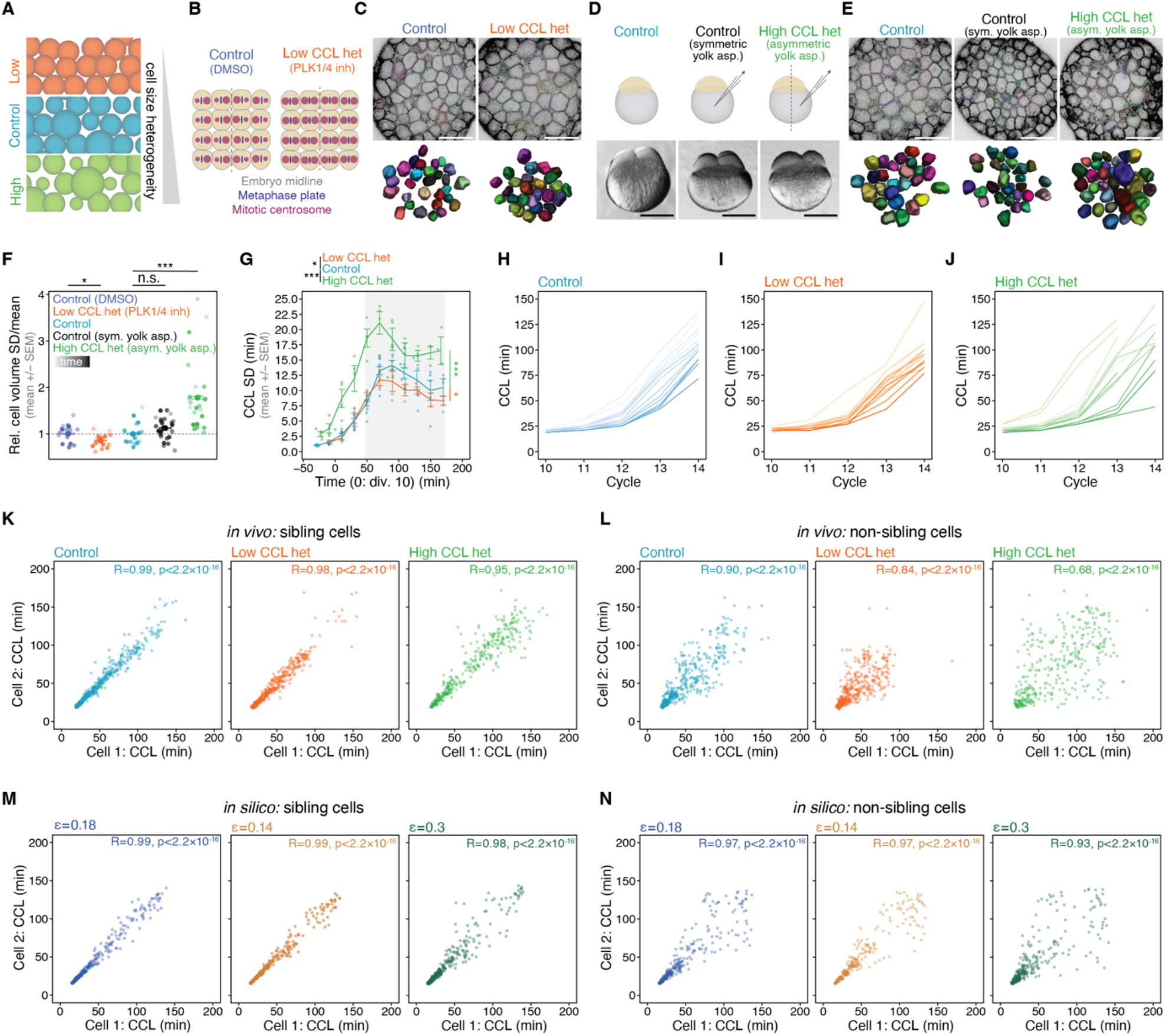
Inherited cell size heterogeneity defines cell cycle length variability. **(A)** Schematic depiction of cell size heterogeneity manipulations. **(B, C)** Low cell size heterogeneity manipulation (PLK 1/4 inhibition). (B) Scheme of the effect of PLK 1/4 inhibition on mitotic centrosome symmetry, adapted from ^84^. (C) Example images (top) of a control (DMSO) embryo and PLK1/4-inhibited embryo and corresponding cell volume segmentations (bottom). **(D, E)** High cell size heterogeneity manipulation. (D) Schematic (top) and example images of embryos with symmetric or asymmetric (high het) aspiration of yolk at 1-cell stage. (E) Example confocal sections (top) of a control (left), a symmetrically (middle) and asymmetrically (right) aspirated embryo and corresponding cell volume segmentations (bottom). (**F)** Variability of cell volume (SD/mean) with low and high CCL het conditions normalized to mean of DMSO-control and control respectively (Control (DMSO): n=503 cells, N=3 embryos; Low CCL het: n=488, N=3; Control: n=460, N=3; Control (sym. yolk asp.): n=798, N=5; High CCL het: n=713, N=4). Opacity indicates time post division 10. Error bars indicate embryo mean +/-SEM. Statistics: two-sided Student’s t-test (low CCL het), two-sided Mann-Whitney-U-test (Control (sym. yolk asp.) and high CCL het (asym. yolk asp.)) **(G)** SD of CCL in cell size heterogeneity-manipulated conditions over time (Control: n=844 cells, N=10 embryos; Low CCL het: n=270, N=3; High CCL het: n=289, N=4). Statistics: Comparison against control (grey box: 50 to 170 min) with two-sided Welch’s test (low CCL het) and two-sided Student’s t-test (high CCL het). **(H-J)** CCL within individual cell lineages throughout CC 10-14 of exemplary embryos in indicated conditions (Control: n=16 cell lineages; Low CCL het: n=15; High CCL het: n=15). Colour indicates condition and colour intensity indicates individual cell lineages. **(K-N)** Pairwise comparison of CCL in sibling and non-sibling cells in indicated experimental conditions from CC 9 onwards (K, L; Control: n=1274 cells, N=3 embryos; Low CCL het: n=1220 cells, N=3 embryos; High CCL het: n=859 cells, N=3 embryos) and from *in silico* simulations at indicated noise ε (M, N; n=50 simulated cells, 100 replicas). Statistics: Spearman Correlation. Asym, Asymmetric; CCL, Cell Cycle Length; Div., Division; Het, heterogeneity; Inh, Inhibitor; SEM, Standard Error of the Mean.; Rel., Relative; Sym, Symmetric; SD, Standard Deviation; Yolk asp., Yolk aspiration. Scale bars: (C, E) 100 µm, (D) 50 µm.

Although both symmetric and asymmetric aspirations have similar mean cell volume (**Fig. S3B**), only the asymmetric aspirations result in blastoderms with cell populations of higher cell volume variability (**Fig. 3F, Fig. S3C**). Despite CCL elongation starting earlier in both symmetric and asymmetric aspirations, due to having fewer resources (**Fig. S3D, E**), only in the asymmetric aspiration we observe a significant increase in CCL variability (**Fig. 3G, Fig. S3F**).

Moreover, we further explored if the above cell size manipulations also affect cell lineage separation. This is based on our finding that the amplification of stochastic differences in resource allocation by the underlying hyperbolic growth of cell cycle elongation is driving cell lineage ranking, suggesting that more similar vs more different cell sizes should lead to less- or more-resolved cell lineages. To this end, we analysed CCLs in cell lineages from control embryos vs embryos of decreased or increased cell size variability, where indeed we observe less-resolved vs more-resolved cell lineages, respectively (**Fig. 3H-J, Fig. S3G-I’**). This is in agreement with the observation that both in experiments and in the model predictions, sibling and non-sibling cells tend to display more similar CCLs in the case of decreased cell size variability (*ε* < 0.18), and less similar CCLs in the case of increased cell size variability (*ε* > 0.18) (**Fig. 3K-N**).

Altogether, the above results pinpoint small but inherited cell size differences as the mechanism driving the large CCL differences through the generations. This raises the question of how this inherited cell size and resource-dependent heterogeneity could impact the tissue material phase transition.

### Temporal CCL variability generates spatial division patterns impacting tissue-level cell-cell contact remodelling

According to the above results, the CCL of a cell relative to the whole cell population is a heritable feature that is driving the de-synchronization process. Spatially, we observe that daughter cells stay rather close to each other until the 13^th^ generation (**Fig. S4A, Movie 3**), raising the hypothesis that inherited CCL temporal differences may generate spatial trends in the division events when observed under chronological time. To explore this hypothesis, we tracked cell divisions and performed a spatial analysis. A pairwise comparison of divisions at each cycle with regard to how closely they happen in time (Δt) and space (Δs) did not show a correlation (**Fig. S4B, C**), in agreement with previous studies showing that large-scale spatial patterns, such as mitotic waves, stop before MBT ^45,86^. To uncover a more subtle relation between the spatial and temporal dimensions, we ranked cell divisions sequentially by the time they take place and computed the spatial distance (Δs) between consecutive divisions (**Fig. 4A, A’**). We then generated a histogram representing the distribution of Δs and compared it to a histogram generated by randomly ranked division times (**Fig. 4B**) ^87^, by computing their distances using the Jensen-Shannon divergence (JSd) ^88^ (**Fig. 4C**). Although the mean JSd between randomly vs sequentially ranked distributions does not show major differences between control, high and low CCL variability conditions, there is a consistent increase in the JSd variability with temporal CCL variability (**Fig. 4D, Fig. S4D**). This suggests that the degree of temporal differences in cell divisions defines how heterogeneous their spatial distribution is within the tissue.

**Figure 4.**
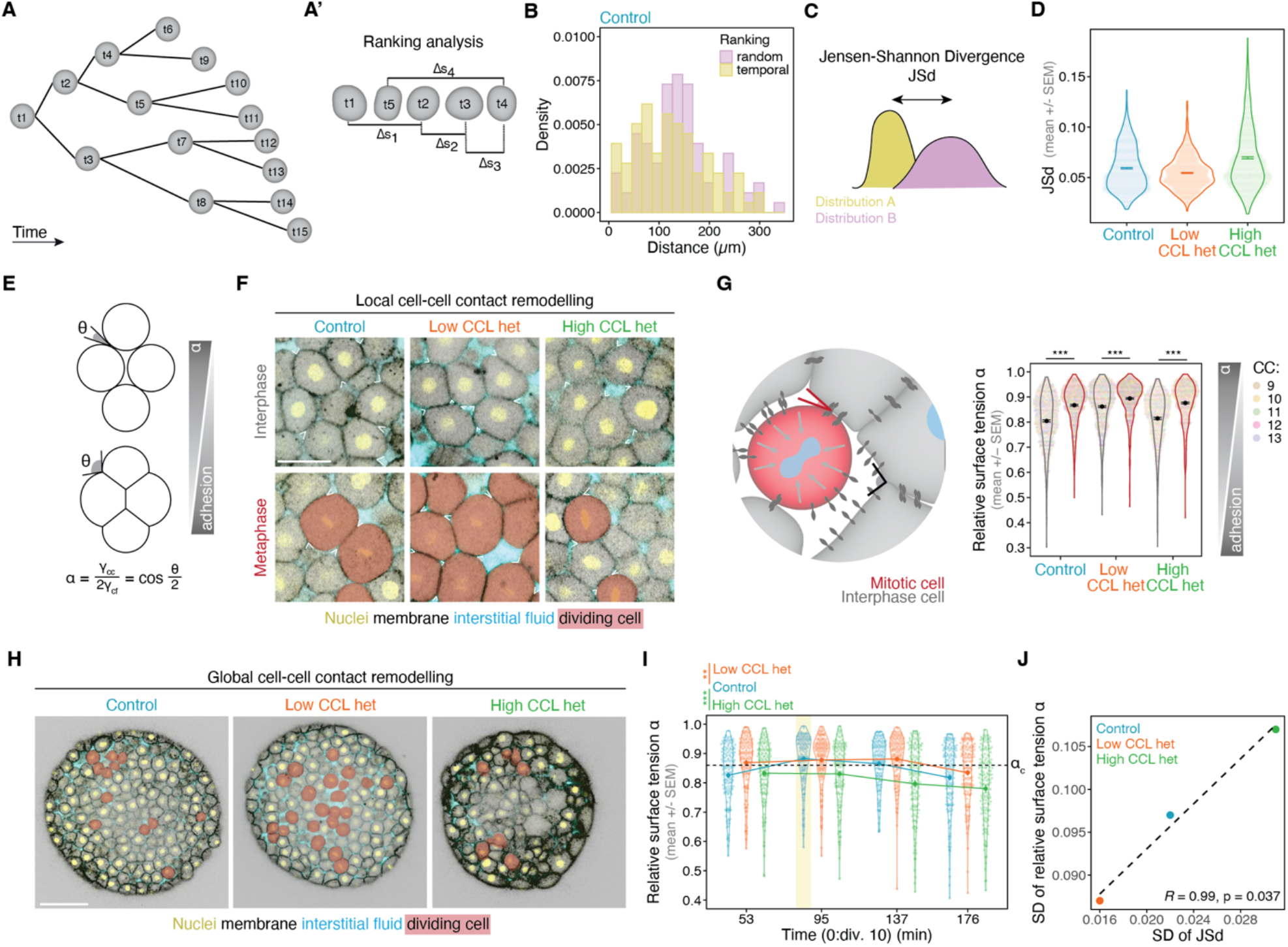
Temporal CCL heterogeneity defines spatial division events tuning global cell-cell contact remodelling. **(A, A’)** Schematic depiction of temporally ranked cell division analysis: cell divisions are ranked in time and the spatial distance Δs is computed between consecutive division events (temporal ranking). As control, the divisions are also ranked randomly (random ranking) and the Δs between random divisions is calculated. **(B)** Example histogram of Δs computed from randomly or temporally ranked division events in a control embryo (n=87 divisions). Only cell divisions around the onset of tissue fluidization are considered (temporal window 50-100 min post div. 10) for this analysis. **(C, D)** To assess the difference between the distribution of Δs in the random and temporal ranking, the Jensen-Shannon Divergence (JSd) is calculated for 300 randomisations per embryo (N=3 embryos per condition). **(E)** Schematic depiction of the effect of mitotic rounding on cell-cell junctions and cell-cell adhesion after the Young-Dupré relationship (bottom). **(F)** Example images of cells in interphase (top) or metaphase (bottom) CC 11 in indicated experimental conditions with dividing cells highlighted in red. **(G)** Relative surface tension α computed from angle measurements in different cell size manipulation conditions (Control: n = 1323 angles, N=3 embryos; Low CCL het: n=1300, N=3 embryos; High CCL het: n=1100, N=3). Statistics: two-sided Mann-Whitney-U-test between interphase and metaphase in each condition. **(H)** Example images of embryos during CC 12 in indicated conditions with dividing cells highlighted in red. **(I)** Relative surface tension α computed from angles measured at the indicated time points post division 10 (control: n=600 angles, N=3 embryos; Low CCL het: n=600, N=3; High CCL het: n=591, N=3). Statistics: two-sided Mann-Whitney-U-test. **(J)** Correlation of SD of relative surface tension α and of SD of JSd between random and temporal rankings of cell divisions during 50-100 min post division 10 (N=3 embryos per condition, with n=300 randomisations for JSd values per embryo. Angle measurements (surface tension α) per condition were: Control: n=300 angles; Low CCL het: n=300; High CCL het: n=291). Statistics: one-sided Pearson Correlation. Dashed line: linear fit. CC, Cell Cycle; CCL, Cell Cycle Length; Div., Division; Het, heterogeneity; JSd, Jensen-Shannon Divergence; SEM, Standard Error of the Mean; SD, Standard Deviation. Scale bars: (F) 50 µm, (H) 100 µm.

Previous work showed that during mitosis cell-cell contacts remodel ^23,24,26,27^, raising the question whether the amount of CCL variability defines the spatiotemporal patterns of cell-cell contact remodelling within a tissue. To this end, we examined local vs global effects of mitosis-mediated cell-cell contact remodelling under different levels of CCL variability. To do this, we quantified the cell contact surface tensions during the cell cycle, a cell physical property associated with cell-cell adhesion strength ^89,90^. This can be derived from the Young-Dupré relation, where a non-dimensional parameter *α*, corresponding to the ratio of cell-cell and cell-extracellular fluid surface tensions, is experimentally obtained from the angle at the cell-cell contact edge (**Fig. 4E**, see Supplementary Theory Note) ^91^. Quantifying relative cell surface tension during division showed that each dividing cell locally lowers its adhesion to its neighbours (higher *α*) when compared to interphase cells (lower *α*). This implies that the capacity of the mitotic rounding to locally reduce cell-cell adhesion strength is retained in the experimental conditions where CCL variability is changed (**Fig. 4F, G**). Importantly, at the tissue-scale it has been previously shown that the relative surface tension *α* acts as a control parameter for a rigidity percolation transition by affecting the global cell connectivity patterns ^8^. By increasing *α*, a large number of connections is lost at a critical point of *α, α*_*c*_ ≈ 0.866, triggering the cell connectivity to cross the isostatic critical point for generic rigidity *k*_*c*_, and induce tissue fluidization ^8^ (**Fig. S4E**). Given that CCL variability changes the spatial distribution of dividing cells, one would expect that, within a wider cell neighbourhood, the degree of co-occurrence of dividing cells will affect the tissue-level distribution of contact surface tensions, and presumably the rigidity transition (**Fig. 4H**). To this end, we quantified relative surface tension *α* at the global tissue-scale, including both interphase and metaphase cells. We observe that control embryos show a slight increase in *α* at the 13^th^ generation (∼100 min post-MBT, corresponding to the timing of tissue fluidization). Embryos with low CCL variability have overall slightly higher mean and less variable *α* values throughout, and embryos with high CCL variability have overall lower mean and more variable *α* values (**Fig. 4I, J, Fig. S4F**). Strikingly however, only control embryos, which exhibit intermediate levels of CCL variability, cross *α*_*c*_ exactly at the 13^th^ generation (**Fig. 4I**, yellow band). This result shows that, although each division event can locally remodel the immediate adhesion to its direct neighbour (**Fig. 4G**), at the tissue-level, CCL variability defines the spatial variability in relative surface tension *α* within the tissue (**Fig. 4J**) in such a way that at a certain CCL variability amount, the tissue crosses the critical point in *α* which triggers the rigidity transition.

Overall, the quantitative link between temporal, spatial and rheological dimensions suggests that stochasticity levels define a regime of CCL variability in which the right number of cell-cell contacts is simultaneously remodelled. This level of remodelling would allow tissue-wide fluidization at a specific and reproducible time, marking the onset of morphogenesis. These findings imply that an optimum variability regime may exist at which the material phase transition is robust, meaning that it can tolerate microscopic disorder.

### An optimum level of CCL variability times and tunes tissue-level mechanical transitions at the onset of morphogenesis

To address whether the rigidity transition occurs robustly and timely only when an optimum level of CCL variability is reached, we sought to tune the latter without affecting developmental rate and quantified blastoderm rigidity, both *in silico* and *in vivo*. We designed a mechanical, 3D realistic model that allows active contact remodelling and cell duplication, and explored tissue rigidity at different levels of CCL variability, taking into account the cell cycle-dependent dynamics of the cell mechanical properties. In particular, we developed a 3D centre-based cell model that represents cells as foam bubbles with surface tension and adhesive tension, as determined by the Young-Dupré relation (**Fig. 5A**). We assumed isovolumetric cell divisions in random directions such that the tissue has constant volume, as expected for the cleavage period ^48^. We introduced the CCL evolution governing cell division times from the stochastic dynamical equation derived above, equation (2), using the parameters obtained from the fittings of the CCL and variability, and simulated mitotic rounding based on the tension balance at the cell-cell contacts (**Fig. S5A**). We used the experimentally measured *α* values for mitotic rounding and defined the rate of cell rounding as proportional to the CCL (**Fig. S5B**). By tuning the noise term *ε* from equation (2) we can incorporate the level of CCL variability to match the crossing of generations and synchronicity observed in control embryos (*ε* ≈ 0.18) (**Fig. 5B**), but also the experimental conditions of low (*ε* < 0.18) vs high (*ε* > 0.18) CCL variability. From the large-scale 3D simulations, we performed rigidity percolation analysis on tissue slices (**Movie 4, Fig. S5C**). Remarkably, we observe that, at *ε* ≈ 0.18, the noise level corresponding to control embryos, we capture the dynamics of the physiological tissue rigidity transition, where the critical point in connectivity is reached only at the peak of the 13^th^ generation (**Fig. 5C-E**), matching exactly the timing of the fluidization transition in control embryos (**Fig. 5F-H, Fig. S5D-G**).

**Figure 5:**
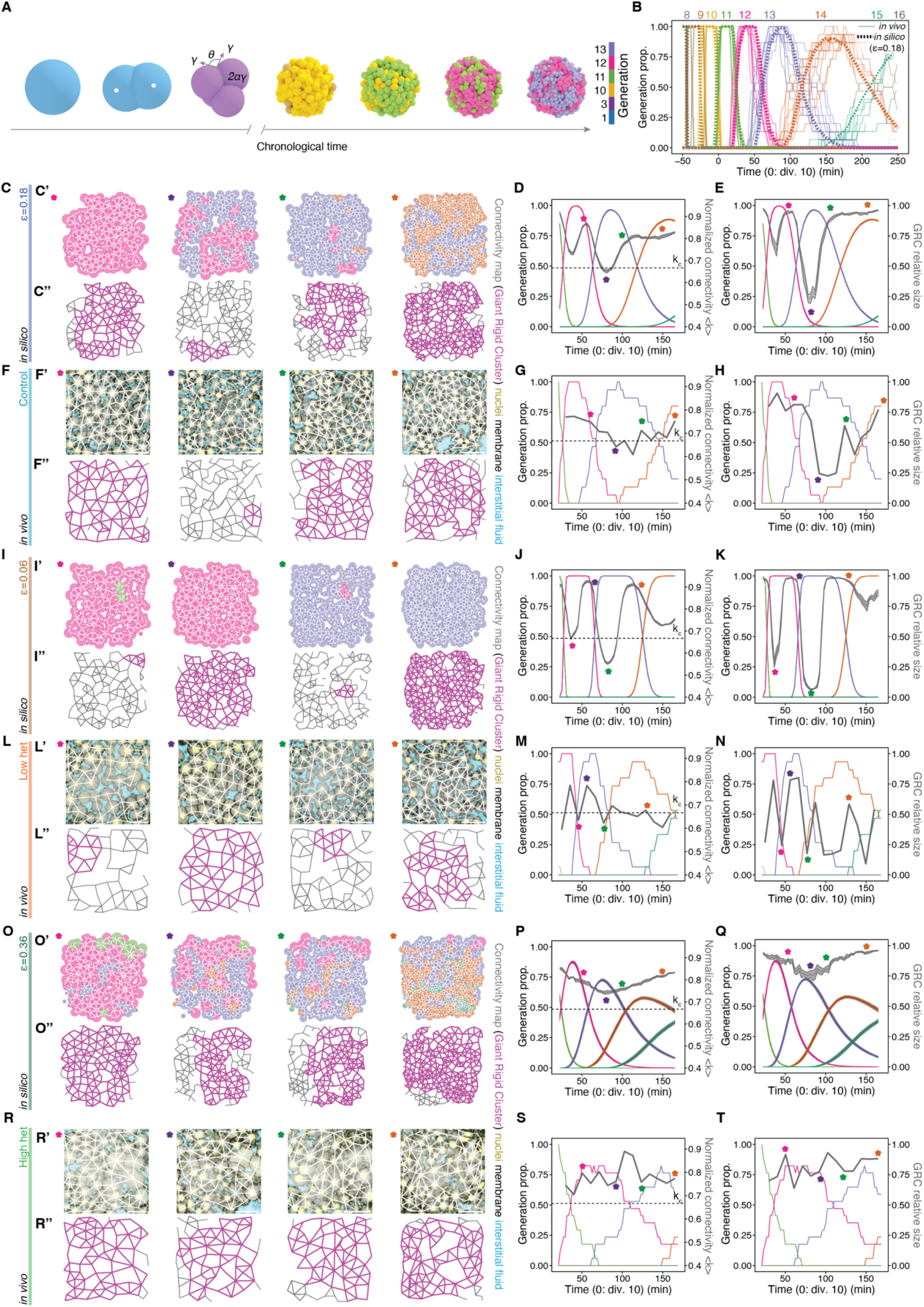
The level of cell cycle length heterogeneity defines the dynamics of tissue fluidization. **(A)** Schematic depiction of a 3D centre-based cell model representing cells as foam bubbles. Colour indicates cell generation. **(B)** Correlation of generations resulting from the simulation (*in silico ε*=0.18; dashed lines) and of *in vivo* experiments over time (same experimental data as in Figure 1F-G and S1D; n=920 cells, N=10 embryos). **(C-T**) Exemplary networks and cell generations overlaid with network parameters over time (average connectivity <k> and GRC size) in cell cycle heterogeneity-manipulated *in silico* (C, I and O) and *in vivo* (F, L and R) tissues: controls (C-H; simulation *ε* = 0.18), low CCL het (I-N; simulation *ε* = 0.06) and high CCL het (K-N; ε = 0.36). *In silico* data is shown as mean +/-SEM of 25 simulations of 35 networks per *ε. In vivo* data shows individual embryos (Control: n=14 networks and 57 cells for generations; low CCL het: n=14 networks and 53 cells for generations; high CCL het: n=14 networks and 52 cells for generations). Asterisks at networks indicate timing in graphs on the right. CC, Cell Cycle; CCL, Cell Cycle Length; Div., Division; Generation prop., Generation proportion; GRC, Giant Rigid Cluster, Het, heterogeneity; SEM, Standard Error of the Mean. Scale bars: (F’, L’, R’) 50 µm.

Consistently, by reducing CCL variability in the simulations, corresponding to noise levels in the stochastic differential equation *ε* < 0.18, we observe a “flip-flop” pattern of repeated fluidization events (**Fig. 5I-K**), where the tissue crosses the connectivity critical point <*k*_*c*_> earlier and multiple times. Remarkably, performing a tissue rigidity analysis in PLK1/4 inhibited embryos, which have low CCL variability, showed the exact same phenotype, where the embryo undergoes repetitive rounds of fluidization-rigidification (**Fig. 5L-N, Fig. S5H-K**), demonstrating that low CCL variability leads to an earlier transition and unstable tissue mechanical states. Conversely, when increasing variability in the simulations (*ε* > 0.18), we observe that the tissue does not reach the critical point in connectivity and does not sufficiently fluidize (**Fig. 5O-Q**). Strikingly, this phenotype matches the tissue rigidity patterns of the asymmetrically aspirated tissues, where they mostly fail to cross the critical point of the rigidity transition and therefore stay rigid (**Fig. 5R-T, Fig. S5L-R**).

The combined *in silico* and *in vivo* results shown above demonstrate that the onset of embryo morphogenesis occurs at an intermediate level of biological noise, strongly supporting the presence of an optimum regime at which tissue mechanical changes may be most robust. To explore this, we simulated tissue mechanical states along a larger range of noise levels (from *ε* = 0 to *ε* = 0.36) (**Movie 5**) and quantified how many times the tissue crosses the connectivity critical point, as a signature of mechanical stability. We detect an overall inverse relationship between CCL variability and tissue mechanical stability both *in silico* and *in vivo*, with low CCL variability leading to multiple crossings of the rigidity critical point, and high variability preventing any crossing (**Fig. 6A, B**). Crucially, we identify signatures of mechanical robustness around the experimentally measured CCL variability (from *ε* = 0.1 to *ε* = 0.2). Within this range of CCL variability, the tissue mechanical behavior is unaffected despite varying noise levels (**Fig. 6A**, blue shaded area) and corresponds to the highest probability for a single round of a rigidity transition (**Fig. 6C**, blue shaded area). Finally, along this large range of noise levels, we further observe a positive correlation between CCL variability and the timing of the tissue mechanical transition both *in silico* and *in vivo*, with low CCL variability leading to an earlier transition and high CCL variability to later or no transitions (**Fig. 6D, E**). Crucially, we identify signatures of temporal robustness, where around the experimentally measured CCL variability (from *ε* = 0.1 to *ε* = 0.2), the timing of the transition is preserved (**Fig. 6D**, blue shaded area), showing that within a range of cellular heterogeneity, the mechanical transition occurs at the right developmental stage, timing the onset of embryo morphogenesis.

**Figure 6.**
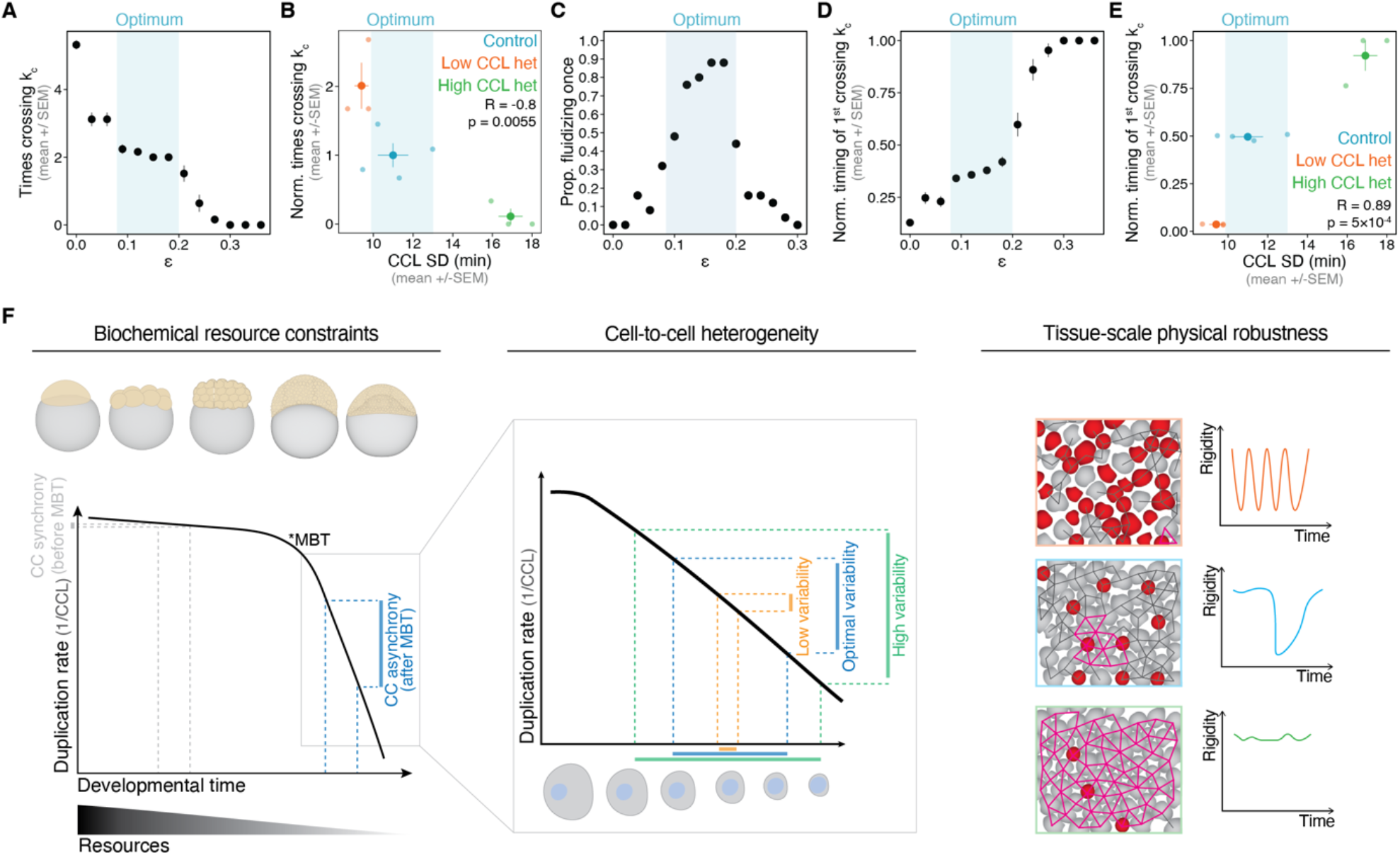
An optimum level of CCL variability times and tunes the rigidity transition at the onset of embryo morphogenesis. **(A)** Number of crossings of the critical point in simulations at the indicated *ε* (N=25 simulations, mean +/-SEM). Blue background indicates robust regime, where the critical point k_c_ is crossed twice first fluidizing and then solidifying the *in silico* tissue. **(B)** Correlation of number of crossings of k_c_ and CCL SD in embryos normalised to the mean of the control condition. **(C)** Probability of fluidizing once (crossing k_c_ twice) of simulations at the indicated *ε*. **(D, E)** Timing of the first crossing of k_c_ normalised to the observation period in simulations (D) and experiments (E). (B, E) Statistics: Pearson correlation; control: n=52 networks and 230 cells for CCL data, N=4 embryos; Low CCL het: n=42 networks and 191 cells, N=3; High CCL het: n=40 networks and 148 cells, N=3. (A, C, D) N=25 simulations of 35 networks per *ε*. **(F)** Schematic summary of the effect of CC heterogeneity on the tissue phase transition at the onset of morphogenesis. Cell size affects resource availability throughout development and within each stage. After MBT, availability of resources limits the cell duplication rate of differently sized cells causing asynchrony under chronological time (left). The degree of cell size heterogeneity then determines the CCL heterogeneity and cell division synchrony (middle). This in turn controls local remodelling of cell-cell adhesion through mitotic rounding (red cells), which globally affects tissue rigidity dynamics (right). A synchronously dividing tissue undergoes multiple fluidization-rigidification cycles (top), while a very asynchronous tissue cannot remodel enough contacts to fluidize (bottom). Intermediate CC heterogeneity allows a transition from high to low rigidity and back exactly once, enabling robust development (middle). Statistics: B and E: two-sided Pearson correlation. CC, Cell Cycle; CCL, Cell Cycle Length; GRC, Giant Rigid Cluster; MBT, Mid-Blastula Transition; SEM, Standard Error of the Mean; SD, Standard Deviation.

As a result, CCL variability is self-tuned within an optimum level driving a timely and correct initiation of tissue-scale morphogenesis.

## DISCUSSION

By uncovering the interplay between deterministic and stochastic elements of cell cycle control, this work demonstrates that an optimum level of cell-to-cell heterogeneity in cell cycle duration is timing and tuning rigidity transitions in embryonic tissues. We identify cellular timing heterogeneity both as a source of mechanical robustness but also as a population-level clock for the onset of embryo morphogenesis.

During embryogenesis, the spatial and temporal patterns of tissue material phase transitions are defined by developmental programs directly tuning the dynamics of cellular properties acting as phase transition control parameters ^4^. Here, we show that stochasticity in such properties provides an additional control axis, where spatial variability in contact remodelling regulates the tissue material state. We find that CCL heterogeneity controls tissue connectivity in a cell size-dependent manner; however, the mechanism of this regulation does not arise from size polydispersity ^92^. This is supported by simulations (**Fig. 5**) where CCL variability was tuned in a cell size-independent manner, and still resulted in similar outcomes in tissue rigidity. Instead, the mechanism by which CCL variability controls the rigidity transition is through precisely varying the spatial distribution of mitotic events within the tissue, which defines the amount and timing of contact remodelling. However, for this mechanism to be achieved, momentary stochasticity is not sufficient, but it has to be history-dependent and inheritable. In this way, cell-to-cell differences in division timing propagate across lineages and create spatial trends in local contact remodelling that are coordinated at the collective level. Strikingly, global contact remodelling occurs robustly within a range of CCL variability, corresponding to ∼12% cell volume stochastic differences, which triggers a single fluidization-rigidification transition precisely at the 13^th^ generation, showing that CCL variability acts as collective-level timer for mechanical transitions.

The identification of an optimal window uncovers how biological stochasticity can fine-tune the macroscopic behaviour of critical systems. In statistical physics, critical points are associated with high susceptibility, such that fluctuations are typically viewed as destabilizing perturbations ^93^. In contrast, our results show that heritable cellular variability acts as a regulatory parameter: it buffers the system against premature or erratic transitions while promoting the coordinated crossing of the critical point, ensuring timely and robust tissue transformation. These findings suggest that noise operates as a robustness mechanism, simultaneously stabilising and timing the phase transition.

Remarkably, the regulation of noise levels does not require feedback architectures associated with robustness ^94-97^. Instead, it emerges from fundamental constraints in cell duplication of early metazoan embryogenesis. Specifically, we have recently shown that the reaction kinetics governing the consumption of finite resources during cell duplication which lead to hyperbolic growth of the CCL are evolutionarily conserved ^44^. By adding a stochastic component in resource allocation this triggers explosive but structured dynamics in CCL variability: under resource abundance, reaction kinetics for cell duplication are very fast, thus small cell-to-cell differences in resource concentration have no impact on CCL, producing synchronized cleavages characteristic of the pre-MBT embryo. As resources become limiting, reaction rates slow down dramatically, leading to the hyperbolic elongation of the CCL (**Fig. 6F**). In this regime, the same cell-to-cell differences in resource allocation have amplifying effects on CCL variability, explaining the asynchrony of the post-MBT regime (**Fig. 6F**). Given that cell size / resources are inheritable parameters, both CCL and its variability propagate along lineages and thereby shape both the temporal and spatial distribution of division events. As a result, these constraints give stochasticity a deterministic role: to regulate tissue rigidity in a highly stereotypic fashion, ultimately timing the onset of embryo morphogenesis (**Fig. 6F**). Intriguingly, cell cleavage desynchronization is a process shared across metazoans ^22,47^, and our recent findings showed that the mathematical singularity of the CCL hyperbolic growth actually corresponds to the onset of gastrulation across species ^44^. These results, together with this work showing that cellular heterogeneity is actively timing mechanical morphogenesis during gastrulation, suggests that a deeply conserved interplay between biochemical resource constraints, cellular stochasticity and tissue mechanics may drive the first collective behaviour of metazoan embryogenesis.

In conclusion, we provide an *in vivo* demonstration of how living materials harness microscopic disorder —both at the spatial and temporal scales— to achieve macroscopic mechanical robustness. Cellular activity, heterogeneity and heritability act as knobs, tuning when and how tissues cross critical points of phase transitions. Beyond the cell cycle, heterogeneity in transcriptional states, metabolic activity, motility, and cell shape is key for morphogenesis of diverse multicellular systems –whether pre-patterned or self-organized, clonal or facultative ^12,14,15,18,31,32,98-108^. The coupling of such multi-component biological noise to deterministic, and even evolutionarily constrained, cellular dynamics gives rise to a wide range of tissue dynamical regimes unique to active matter. In this view, noise in living matter is not simply a by-product of microscopic fluctuations: due to biological inheritance, stochastic temporal fluctuations are transformed into a regulatory dimension through which developing tissues achieve robust morphogenesis.

## ACKNOWLEDGEMENTS

We thank Takashi Hiiragi, Zena Hadjivasiliou and members of the Petridou, Corominas-Murtra and Smeets groups for technical advice, critical discussions and feedback on the manuscript; and the European Molecular Biology Laboratory (EMBL) Advanced Light Microscopy Facility (ALMF), the Data Science Centre, and the zebrafish facility for continuous support. We thank Jlenia Vitale for assisting with embryo aspiration experiments. We thank the Heisenberg lab for providing fish lines and plasmids. We thank the Barcelona Collaboratorium for Modelling and Predictive Biology for hosting N.I.P. and B.C-M. during writing sessions. This work was supported by the Weave project “Tissue material phase transitions and their role in embryo pattern formation” from the Deutsche Forschungsgemeinschaft (DFG, German Research Foundation, 518354236, PE 3800/1-1) to N.I.P. and the Österreichischer Wissenschaftsfonds (FWF, Austrian Science Fund, I6533) to B.C-M. B.C-M. and A.A-T. acknowledge the support of the field of excellence “Complexity of life in basic research and innovation” of the University of Graz. N.I.P. was supported by the European Union’s Horizon Europe research and innovation program (European Research Council Starting Grant “CoRe” 101162743). J.V., T.E.B. and B.S. acknowledge support from the Fonds Wetenschappelijk Onderzoek (FWO, Research Foundation Flanders) under project IDs 1266425N and 11D9921N, and from the Onderzoeksraad (Research Council), KU Leuven under project IDs IDN/25/007 and C14/18/055.

## AUTHOR CONTRIBUTIONS

N.I.P. and B.C-M. designed the research. M.S-J. performed all experiments and analysed the experimental data. J.V. performed the large-scale 3D simulations. A.A-T. and M.S-J. analysed the spatial division events data. B.C-M. developed the mathematical model for cell cycle elongation and variability and J.V., T.B. and B.S. developed the computational model for 3D simulations. J.V. and T.B. analysed the 3D simulation results. N.I.P. and B.C-M. wrote the manuscript with input from all authors.

## DECLARATION OF INTERSTS

The authors declare no competing interests.

## SUPPLEMENTARY FIGURES AND FIGURE LEGENDS

**Figure S1.**
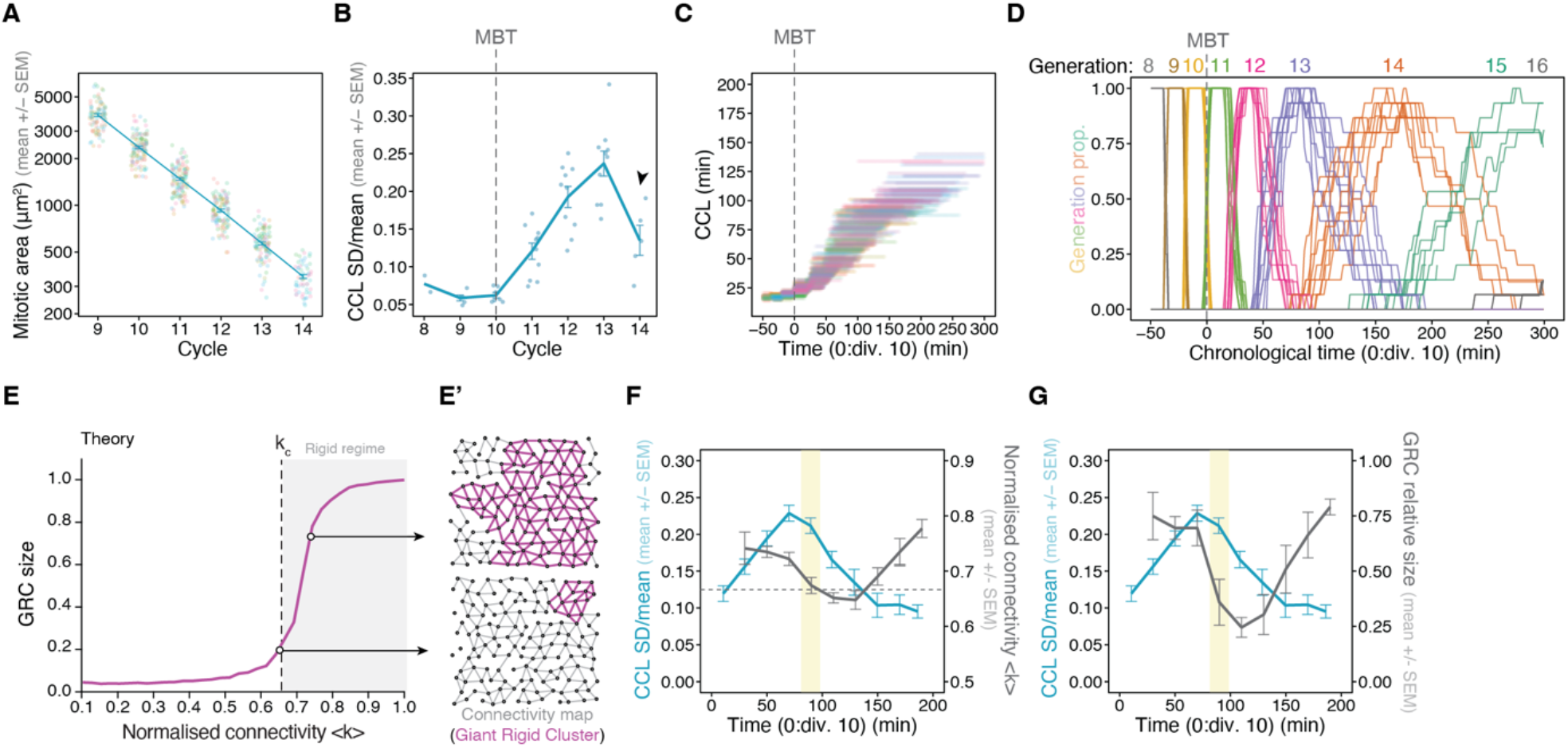
Quantification of cell cycle length and its variability, and tissue rigidity dynamics at the onset of embryo morphogenesis. **(A)** Cell area quantifications at metaphase of indicated cell division (n=657 cells, N=10 embryos, each colour indicates a different embryo). **(B)** Coefficient of variation (SD/mean) of CCL per embryo (n=740 cells, N=10 embryos). Black arrowhead indicates drop of CCL variability. **(C)** Alignment of CCL in chronological time, for which each cell is assigned the length of its current CC until the metaphase plate (n=873 cells, N=10 embryos, each colour indicates a different embryo). **(D)** Generation plot showing the proportion of cells at the indicated generation per embryo over time (each line represents one embryo, n=940 cells, N=10 embryos). **(E)** Plot of the GRC size as a function of network connectivity, reproduced from ^6^. At critical connectivity, k_c_, the GRC size changes non-linearly. (**E’**) Simulated networks with connectivity above (upper panel) or below (lower panel) the critical point, leading to a large vs small GRC, respectively. (**F, G**) Plots of overlaid CCL SD/mean with network connectivity (F) and GRC size (G) (same as in Fig. 1F, G). The horizontal grey dashed lines indicate the connectivity critical point k_c_. CC, Cell Cycle; CCL, Cell Cycle Length; Div., Division; Gen., Generation; GRC, Giant Rigid Cluster; MBT, Mid-Blastula Transition; SEM, Standard Error of the Mean; SD, Standard Deviation.

**Figure S2.**
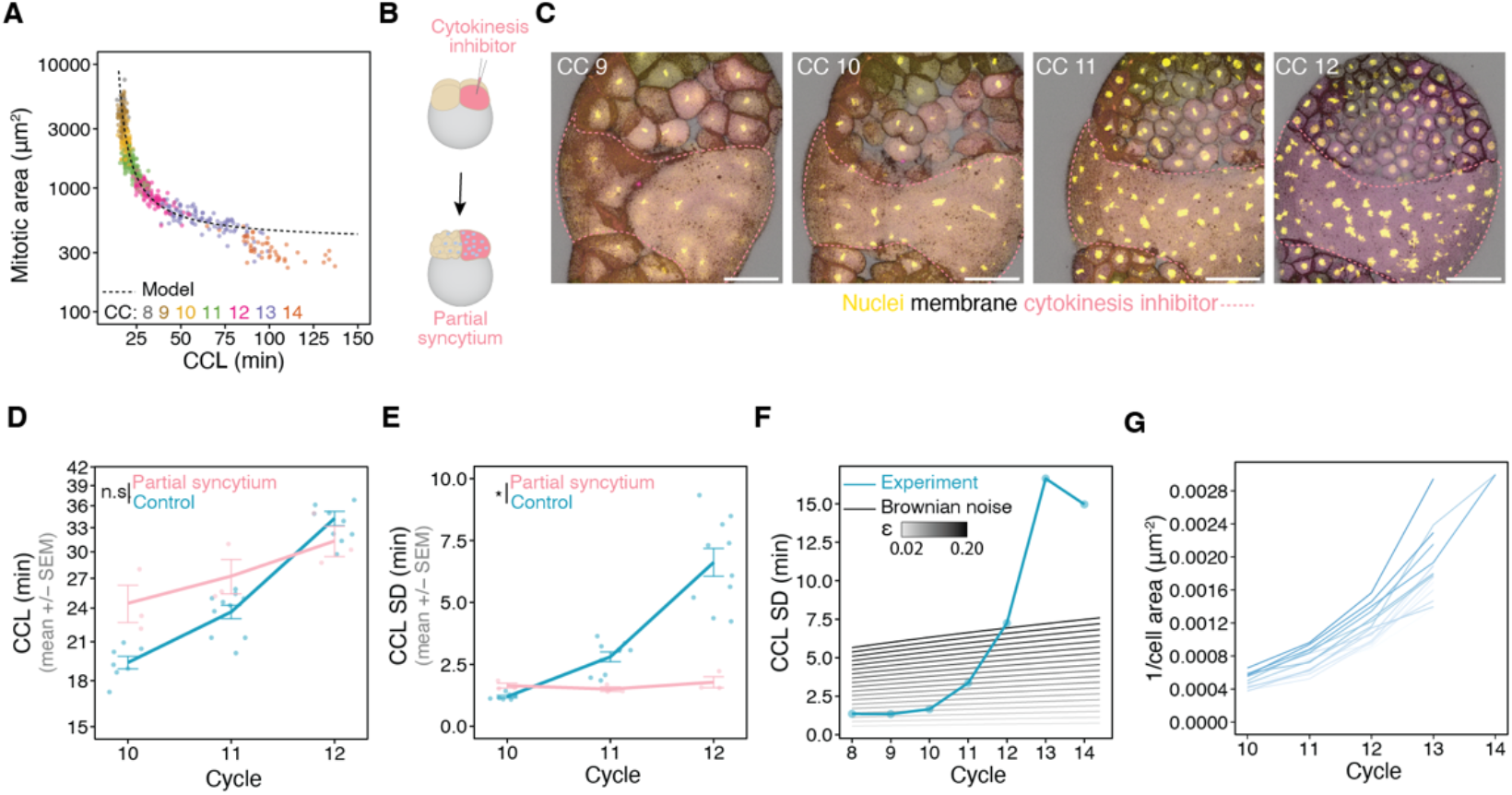
Cell volume differences via cell division control differential resource allocation that in turn defines cell cycle dynamics. **(A)** Correlation of cell size at metaphase and CCL in experimental data (N=8 embryos, n=576 cells) and its theoretical prediction (dashed line). **(B)** Schematic depiction of partial syncytium experiment. Injection of a cytokinesis inhibitor in a single-cell results in a partial syncytial after a few division rounds. **(C)** Exemplary partial syncytium in an embryo (max projections of 6 confocal z-slices) at indicated division formed by co-injection of a cytokinesis inhibitor with dextran-Alexa-647. See also Movie 2. **(D, E)** Mean (D) and SD (E) of CCL in partial syncytia and control embryos (control: n=400 cells, N=10 embryos, same control data as in Fig. 1D, E; Partial syncytium: n=119 nuclear cycles, N=3 embryos). Error bars indicate embryo mean +/-SEM. Statistics: two-sided Mann-Whitney U-test. **(F)** Pooled CCL SD (n=740 cells, N=10 embryos) overlaid with theory predictions of CCL SD considering only Brownian noise at different *ε* values (same experimental data as in Fig. 2E). **(G)** Inverse of cell size over indicated cell cycles measured at metaphase in an exemplary embryo (n=15 lineages). Colour shades indicate individual cell lineages. CC, Cell Cycle; CCL, Cell Cycle Length; Div., Division; SEM, Standard Error of the Mean; SD, Standard Deviation. Scale bars: (C) 100 µm.

**Figure S3:**
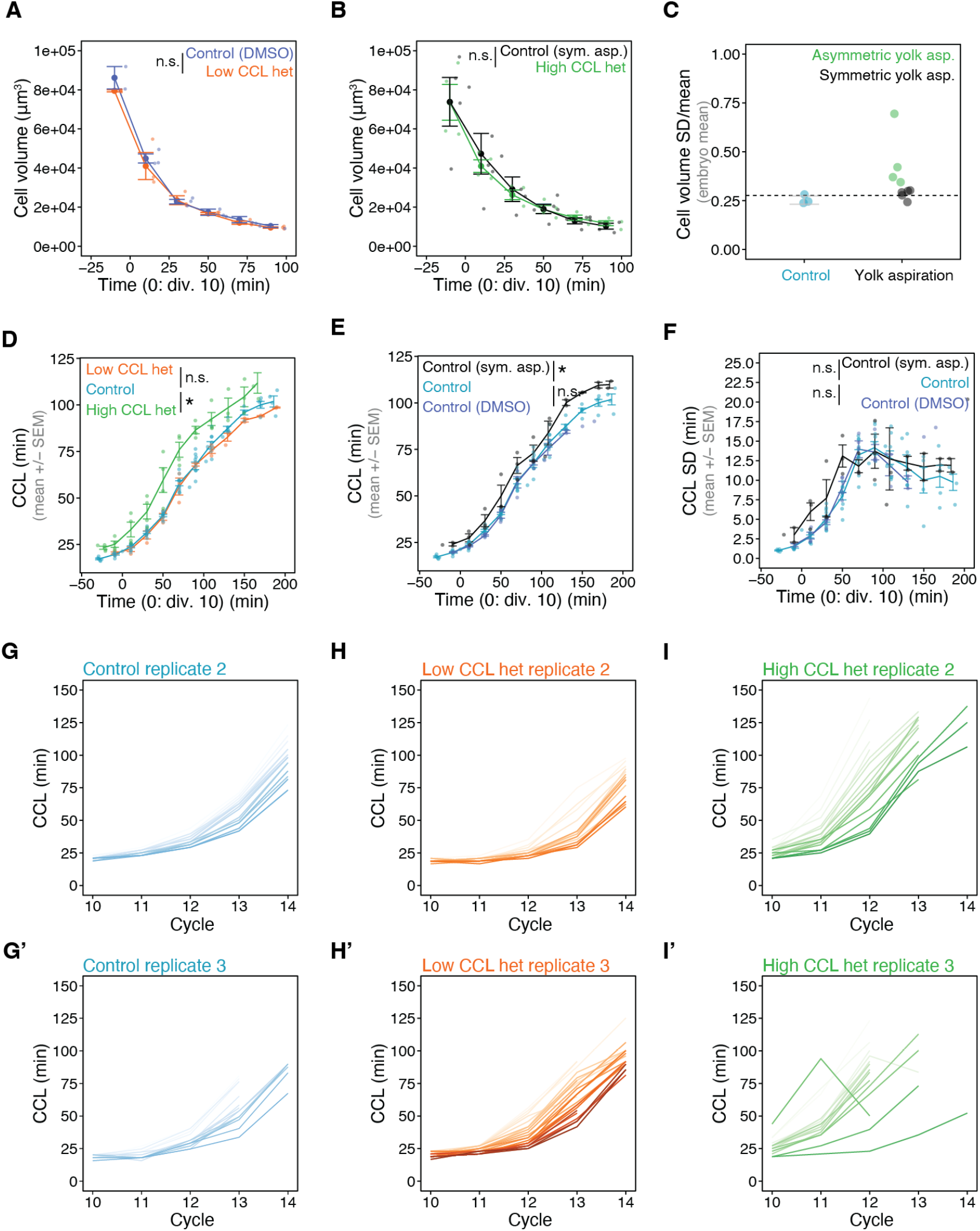
Characterization of cell size manipulated embryos. **(A, B)** Cell volumes over time in experimental manipulations of low CCL het (A) and high CCL het (B). Statistics: two-sided Mann-Whitney U-test (Control (DMSO): n=503 cells, N=3 embryos; Low CCL het: n=488, N=3; Control (sym. yolk asp.): n=798, N=5; High CCL het: n=713, N=4). **(C)** Cell volume variability in yolk aspirated embryos to classify symmetric or asymmetric embryos. Dashed line indicates maximum value of control embryos. **(D)** CCL of indicated experimental conditions over time (Control: n=844 cells, N=10 embryos; Low CCL het: n=270, N=3; High CCL het: n=289, N=4). Statistics: two-sided Mann-Whitney U-test. **(E, F)** CCL (E) and CCL SD (F) of indicated control conditions over time (Control (DMSO): n=295 cells, N=4 embryos; Control: n=844, N=10; Control (sym. yolk asp.): n=201, N=3). Statistics: two-sided Students t-test from 50 to 170 min post division 10. **(G-I)** CCL within individual cell lineages throughout CC 10-14 of exemplary embryos in indicated conditions (G, G’: Control, n=30 cell lineages, N=2 embryos; H, H’: Low CCL het, n=38, N=2; I, I’: High CCL het, n=32, N=2). Colour indicates condition and intensity indicates cell lineage. Asp., Aspiration; CCL, Cell Cycle Length; Div., Division; Het, heterogeneity; Inh, Inhibitor; SEM, Standard Error of the Mean; SD, Standard Deviation; Rel., Relative; Sym, Symmetric.

**Figure S4.**
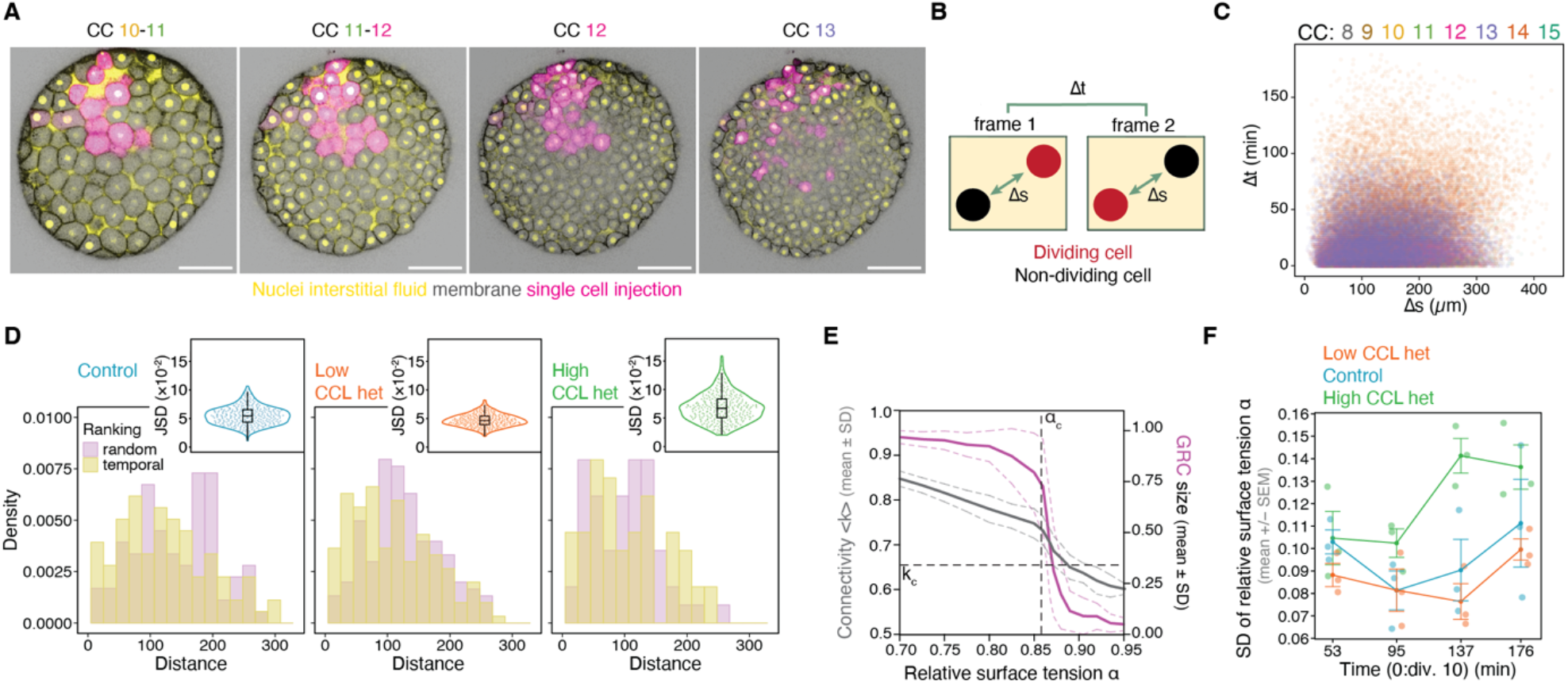
Clonal localisation and analysis of spatial trends in divisions impacting the tissue material state. **(A)** Example of a clonal injection of fluorescently labelled dextran in one cell at the 32-to 64 cell stage. **(B, C)** Schematic depiction of the pairwise comparison between cell divisions (B). The differences in timing (Δt) and spatial distance (Δs) are calculated between all divisions of the same cycle and plotted (C) (n=58618 cell-cell comparisons from 1234 cell divisions, N=3 embryos). **(D)** Exemplary histograms showing distribution of Δs in random and temporal ranking at indicated conditions (Control: n=88 divisions, Low CCL het: n=149, High CCL het: n=56). Several replicas of the random histograms were generated from which a distribution of JSd distances was obtained between the ranked pattern and the random ones (see Supplementary Theory Note for details). Insets show the distribution of JSd values resulting from comparisons between the ranked sequence and 300 randomized sequences, shown separately for each condition for the selected embryo. **(E)** Plot of network connectivity <k>, rigidity (GRC size) and relative surface tension α from *in silico* tissues in which relative surface tension α is decreasing. Critical points (dashed lines) in connectivity and α are marked as k_c_ and α_c_ respectively. Graph modified from ^8^. **(F)** Embryo SD of relative surface tension α from Fig. 4I over time (control: n=1323 angles, N=3 embryos; Low CCL het: n=1300, N=3; High CCL het: n=1100, N=3). CC, Cell Cycle, CCL, Cell Cycle Length; Div., Division; Het, heterogeneity; SD, Standard Deviation. Scale bars: (A) 100 µm.

**Figure S5.**
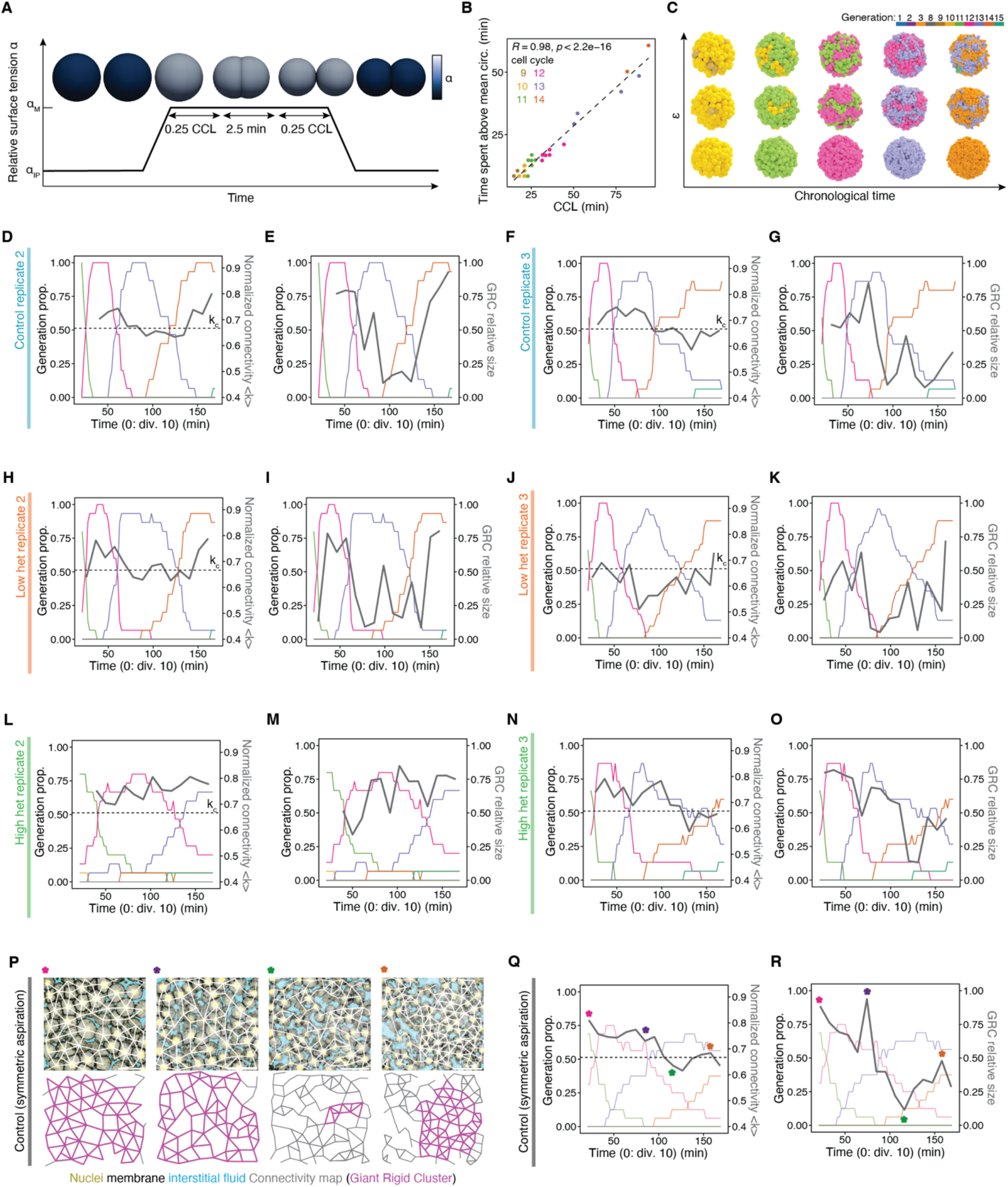
Contact surface tension-driven changes along the CCL and replicates of embryos with altered CCL variability. **(A)** Scheme showing the changes of relative surface tension α throughout a cell division cycle in the 3D centre-based cell model. α_IP_ reflects mean of α measured between cells during interphase and α_M_ reflects the mean of α measured adjacent to mitotic cells (see data in Fig. 4G). In the model, the time duration that cells spend at α_M_ is assumed to be proportional to the CCL, as supported by experimental data (B). **(B)** Correlation of CCL and the time spent rounding, i.e., above the average circularity, for individual cells at the indicated cell cycle (n=37 cells computed from 545 cell area measurements over time, N=2 embryos). Statistics: two-sided Pearson correlation. Dashed line represents linear fit. **(C)** Examples of 3D centre-based cell model simulations over time at varying values of ε. **(D-O)** Connectivity (D, F, H, J, L and N; dashed line indicates critical point k_c_) and GRC size (E, G, I, K, M, O) of exemplary embryos of control (D-G), low (H-K) and high (L-O) cell cycle heterogeneity with generation proportion over time (Control: n=27 networks and 113 cells for generations, N=2 embryos; Low CCL het: n=28 networks and 138 cells for generations, N=2; High CCL het: n=27 networks and 96 cells for generations, N=2). **(P-R)** Exemplary cell networks (P), connectivity (Q) and GRC size (R) of a control (symmetrical aspiration) embryo (n=15 networks and 48 cells for generations). Asterisks at networks indicate timing in graphs on the right. α_IP,_ α at Interphase; α_M,_ α at Metaphase; Cell Cycle Length; Circ., Circularity; Div., Division; Generation prop., Generation proportion; GRC, Giant Rigid Cluster; Het, heterogeneity; SD, Standard Deviation; Sym., symmetrical. Scale bar: (P) 50 µm.

## MOVIE LEGENDS

**Movie 1: Example of cell cleavage rounds in a zebrafish embryo**.

3D view of an example embryo from division 7 onward. Orange: Nuclei and interstitial fluid, grey: cell membrane.

**Movie 2: Embryo with a partial syncytium**.

Max projection of 6 z-slices (4 µm apart) of an embryo with a partial syncytium embryo. Yellow: Nuclei and interstitial fluid, grey: membrane. Top left: time (0: Division 10). Scale bar: 100 µm.

**Movie 3: Clonal dispersion around cell cycle elongation**.

Individual z-plane of an embryo injected with dextran-647 in a single cell at 64-cell stage. Yellow: Nuclei and interstitial fluid, grey: membrane, Top left: time (0: Division 10). Scale bar: 100 µm.

**Movie 4: Example simulations of the 3D-centre based model at varying cell cycle synchrony**.

3D centre-based cell model representing cells as foam bubbles. Cell cycle synchrony was modified by adjusting *ε*.

**Movie 5: Network connectivity and GRC-size changes with cell cycle synchrony in tissue simulations**. Average network connectivity and GRC size over time at varying *εϵ* in simulations. Lines and shaded regions represent mean +/-SEM of 25 simulations of 35 networks per *ε* (from *ε* = 0.00 to 0.36). GRC: Giant rigid cluster.

## MATERIAL AND METHODS

### Fish and embryo handling

Zebrafish (*Danio rerio*) were maintained at 28.5°C under a 14h light / 10 h dark cycle ^109^. The following zebrafish strains were used in this study: Tg(acbt2:H2B-GFP; actb2:Lyn-tdTomato) ^110^, Tg(actb2:Lyn-tdTomato) ^110^ and wildtype A2B2, AB, Golden and TL. Embryos were collected in E3 medium within 5 minutes of laying ensuring synchronisation of each clutch. Embryos were kept at 25°C or room temperature until being imaged at 28°C. Experimental manipulations were performed in Danieau’s medium and staging was performed as previously described ^48^. All animal experiments were conducted according to the guidelines of the Committee for Animal Welfare and Institutional Animal Care and Use (IACUC) under EMBL’s Policy on the Protection and Welfare of Animals Used for Scientific purposes (IACUC code 21-010_HD_NP).

## Microinjections

Unless otherwise indicated, embryos were microinjected at the 1-cell stage with 1nL using a pneumatic pico pump (World Precision Instruments: PV820) with glass capillary needles (30-0020, Harvard Apparatus, MA, USA) prepared with a Flaming/Brown Micropipette Puller (Sutter Instrument CO.: Model P-97). For the labelling of interstitial fluid, embryos were microinjected with 0.5nL of 0.67 μg/μL Dextran-Alexa Fluor 647 (10,000 MW; Invitrogen, Cat. No. D22914) between the cells at 64 to 215-cell stage with a pneumatic pico pump (World Precision Instruments: PV830).

For partial syncytium experiments (**Fig. S2B-F**), embryos were microinjected with 0.5nL of 1mM Aurora kinase inhibitor ZM-447439 (Tocris: 2458) and 0.3 μg/μL Dextran-Alexa Fluor 647 (10,000 MW; Invitrogen, Cat. No. D22914) in 50% DMSO in one cell at the 4-to 32-cell stage. For PLK1/4 inhibition, embryos were microinjected with 1.5μM BI2536 in 1% DMSO in water (Sellekchem: S1109) or 1% DMSO in water, based on ^71^. For lineage visualisation (**Fig. S4A**), embryos were microinjected with 0.5nL of 0.3 μg/μL Dextran-Alexa Fluor 647 in one cell at the 32-to 64 cell stage.

### Yolk aspiration

Yolk material was aspirated close to the cell-yolk interface from one side (asymmetrically) or the centre (symmetrically) from 1-cell stage embryos. Aspiration was performed using a transplantation kit set up consisting of a 1mL BD syringe (SMS medipool: 303172), tubing (VWR International GmBH) and a 6.3mm electrode handle (World Precision Instruments: 2505) with a glass capillary needle prepared as described above. Cell size (a-)symmetry was assessed at the 2-to 4-cell stage and embryos were selected as symmetric or asymmetric. Final assessment of cell size heterogeneity was performed through cell volume segmentation and aspirations classified as symmetric or asymmetric based on that (**Fig S3C**).

### Image acquisition

Dechorionated embryos were embedded with the blastoderm facing upwards in 0.6% low melting agarose within a 2% agarose mold and imaged with a W Plan-Apochromat 20x/1.0 Corr DIC M27 75mm objective on an upright confocal microscope (ZEISS LSM 980) within an incubation chamber at 28.5°C. Images were acquired in ZEN3.3 (blue edition) software (Carl Zeiss). Each embryo was imaged with 40 planes (z-spacing 4μm) of 1024 × 1024 pixels at 8-bit roughly every 2 min with acquisition of lasers 488nm, 561nm and 639nm on one track. Scanning was bidirectional at a speed of 0.77μm/s. Brightfield images were acquired on a Leica M205 FCA stereoscope equipped with a Leica DFC 7000T camera.

### Data analysis and quantification

Image analysis was performed in Fiji ^111-113^ unless indicated otherwise. Except for rigidity analysis, data analysis was done in R studio ^114,115^ with the packages dplyr ^116^ and tidyverse ^115^.

#### Data preparation

For live imaging, different acquisitions of each embryo were manually aligned in z over time and the time interval computed as the weighted average. Embryos were staged based on their cell size at metaphase over at least 3 divisions (see **Fig. S1A**). For staging of yolk aspirated embryos with decreased cell size, cell number in field of view was considered and staging manually adjusted accordingly. The provided movies were stabilised according to the histone channel using the built-in drift correction plugin in Fiji. **Movie 1** was created in Imaris.

#### Cell tracking

Cells were manually tracked using the TrackMate plugin in Fiji ^111-113^ by marking cell divisions (metaphase plate). Cell cycle length was calculated to be the temporal difference between consecutive metaphase plates within one track. 15 cells were followed without siblings in each embryo for calculation of average and variance of cell cycle lengths. For sibling cell comparisons, full lineage trees were constructed of each embryo of at least 15 cells from division 9 onwards.

#### Dynamic analysis of cell cycle length and generation plotting

To analyse cell cycle dynamics in real time, each embryo was aligned by its average time of the 10^th^ division, as defined by tracking and their cell size staging. Each cell was then assigned its cell cycle length for its entire cell cycle duration since its previous division (see **Fig. S1C**). The cell cycle length average, standard deviation and coefficient of variation (SD/mean) were calculated at each time point from at least 15 cells per embryo, except in the syncytium condition, where in one embryo only 10 cells were followed. The average of each embryo within bins of 20-min intervals was calculated and plotted over time. For generation plots, the same calculation was performed based on which cell cycle each cell was in at each time point. For this, the n^th^ 1 division is considered to end the n^th^ cell cycle/generation and cells progress to the next cell cycle/generation i.e., cell cycle 1 and generation 1 are the 1-cell stage and end with the 1^st^ division.

#### Cell size measurements

Cell area was measured by circling the widest cross-sectional area of each cell at metaphase using the freehand tool in Fiji ^111^. Cell size at metaphase was correlated with cell cycle length of the previous cycle leading up to the metaphase (**Fig. S2A**). Cell volumes for cell size variability analysis were measured with the LimeSeg plugin in Fiji using manually placed circles at the cell centre as seeds ^117^. For sibling cell size comparison, volumes of both daughter cells were measured following formation of the nuclear envelope post cell division. Their size ratio was calculated by dividing the larger by the smaller sibling cell volume (**Fig. 2H**). All segmented volumes were manually checked.

#### Circularity measurements

Circularity was measured by circling the widest cross-sectional area of each cell throughout the cell cycle roughly every 2 minutes using the freehand tool in Fiji ^111^. The mean circularity per cycle was computed for each cell and calculated how much time (frames x imaging interval) was spent above it (**Fig. S5B**).

#### Cell network analysis

Cell network analysis was based on previously published work using embryos with labelled interstitial fluid ^8,118^. Cell connectivity networks were constructed from 2D confocal sections of the first to second deep cell layer. Interstitial fluid was binarized using the isoData threshold and smoothed using a 1px median filter in Fiji or binarized manually. Cell segmentation was done using cellpose v2 or v3 ^119^ and corrected manually. Cell segmentation was then used to create cell connectivity networks that were manually corrected using a custom-made Fiji plug-in ^8^. Rigidity analysis of cell networks was based on the pebble game algorithm ^6,50^ and performed as described in ^8^ using a custom-made python script (Python 3.9.14, PyQt5 5.15.9) in Spyder (5.4.5) on macOS with the packages numpy (1.25.2) ^120^ and pandas (2.0.3). For both experiments and simulations, only networks with at least 50 cells, about 20 minutes post division 10, were considered for the analysis, due to size effects in small networks ^6^. All connectivity values, <k>, were normalized to 6, the maximum number of contacts in a 2D network. For real time analysis, networks were aligned based on the 10^th^ division in the embryo or tissue simulations. For **Fig. 1**, networks were pooled and averaged within bins of 20 minutes over the aligned time.

For robustness quantifications (**Fig. 6**), only cell cycle and network data spanning 20-165 min post division 10 were considered for cell size-manipulated and control embryos as well as tissue simulations. Crossing of the critical point was counted when the interval of <k> between one network and the next network 10 minutes later included k_c_ (**Fig. 6A-B**). For simulations, the same analysis was performed every 2.8 minutes (**Fig. 6A**). Crossing events in the experimental data were normalized to the mean of the control (**Fig. 6B**). The timing of the first crossing of k_c_. was normalized to the observation period (0: 20 min and 1: 165 min) and 1 assigned if k_c_. was never crossed (**Fig. 6D-E**).

#### Angle measurements

Angles were measured manually in Fiji ^111^. For the measurements per cell cycle, angles were measured between cell pairs in interphase or at least one cell in metaphase on the plane of the widest angle of the cell-cell contact.

#### Pairwise comparison of cell division timing

For **Fig. S4C**, all cell divisions were compared pairwise in each generation per embryo and the difference between their timing (Δt) and spatial location (Δs) were calculated. Δs was calculated in 3D as follows:

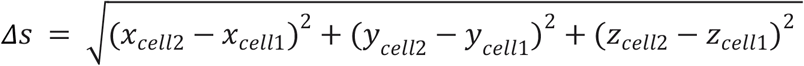

### Cell cycle length comparison of sibling and non-sibling cells

Cell cycle length of sibling cells was compared pairwise within each generation in each embryo (**Fig. 3K-N**). In the experimental data, all non-sibling cells were compared within each generation and a sample of the same number as sibling cell comparisons was taken per generation from each embryo to compare equal sample size between sibling and non-sibling cells. For *in silico* data, a sample of the same number as sibling cell comparison was taken from non-sibling cell comparisons regardless of the generation.

### Spatiotemporal similarity analysis using Jensen–Shannon divergence

To quantify the relationship between the temporal ordering of cell divisions and their spatial organization (**Fig. 4, S4**), we measured the similarity between spatial distance distributions using the Jensen–Shannon divergence (JSd). For each embryo and experimental condition, cell divisions occurring within the 50–100 min time window were ranked sequentially according to their division timing (Δt). Spatial distances (Δs) were then computed between consecutive divisions in this temporally ordered ranking, generating a distribution of Δs values associated with temporally close cell divisions (ranked condition). To generate a null-model distribution in which temporal and spatial relationships are uncoupled, the same set of division events was randomly permuted, and Δs values were recalculated between consecutive events in the randomized ranking (random condition). Histograms of Δs values were constructed for both ranked and random conditions using identical binning and normalization.

The similarity between the ranked and random Δs distributions was quantified using the Jensen–Shannon divergence ^88^. For each embryo and condition, the randomization procedure was repeated 300 times, generating multiple randomized Δs histograms. Each randomized histogram was compared to the ranked histogram, yielding a distribution of JSd values that captures the expected variability due to stochastic sampling. These distributions are summarized as violin plots per condition in **Fig. 4** and **S4**. To study the spatiotemporal organization to tissue material properties, we additionally compared, across embryos, the standard deviation of the JSd distributions with the standard deviation of the material parameter α (**Fig. S4J**).

### Plotting and fitting

Plots were generated using ggplot2 in R studio with the packages ggplot2 (3.5.1) ^114^, dplyr (1.1.4) ^116^, ggh4x (0.3.1), ggpubr (0.6.2), scales (1.3.0) and RcolorBrewer (1.1.3).

For **Fig. 1**, to show that the data do not follow an exponential trend, we plot, as a visual reference, different exponential curves crossing the experimental data at generation 8:

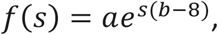

where *α*=16.86 and *b* was chosen as 0.01, 0.1, 0.2, 0.3 or 0.4 for the different trends.

#### Statistics

Statistical analysis was performed in R (version 4.4.1) with the packages ggplot2 (3.5.1) ^114^, ggpubr (0.6.2) ^121^, dplyr(1.1.4) ^116^, tidyr (1.3.1), tidyverse (2.0.0) ^115^, gtable (0.3.5), broom (1.0.6) and car (3.1.3). Statistical tests and sample sizes are provided in the figure legend. Data was checked for normality with the Shapiro-Wilk test and for homogeneity in variances with the Levene’s test. For two-group comparisons, a two-tailed Student’s t-test was used for parametric distributions with equal variance, Welch’s t-test for unequal variance and Mann-Whitney U-test for non-parametric distributions. Correlations were calculated with the Pearson or Spearman method as indicated in figure legends.

## Data availability

All data generated in this study will be publicly available upon manuscript publication.

## Code availability

All codes are available in the following GitHub repositories:

Rigidity analysis: https://github.com/coldlaugh/pebble-game-algorithm/blob/master/pebble.pyx

Fiji plugin for connectivity network reconstruction: https://github.com/Petridou-Laboratory/cell-connectivity-networks

## I. DETERMINISTIC DYNAMICS

Here we consider only the cell cycle length and we omit heterogeneity in the volumes. This can be seen either as 1/a mean-field approximation of the cell-cycle length evolution or 2/ a syncytium, where all the cells share the same resources. The reasoning for this deterministic approximation is taken from reference [1].

### A. Enzyme kinetics and cell-cycle elongation

Given a substrate concentration *c*, a maximum reaction rate *u*_*M*_ and a kinetic constant *K*_*m*_, the equation of Michaelis-Menten for the reaction rate *u* of a given enzyme as a function of the underlying substrate concentration reads:

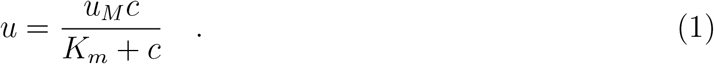

This equation tells us how much product is generated per time unit. For low values of *c* it has a linear behaviour:

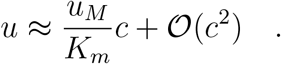

For the first division round, we have an initial concentration of *c*_0_ = *N*_0_*/V* of the substrate, being *N*_0_ the actual amount of substrate molecules and *V* the volume of the first cell. The overall volume is considered invariant throughout the cleavage period. For division *s* we assume that resources are being consumed at a certain pace 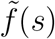, being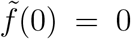 –no resource has been consumed at the starting point– and 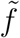 is a monotonously increasing, positive function of *s* and, for simplicity, we assume that 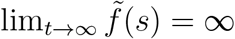 and that 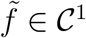. In consequence, the concentration after the first *s* division rounds, *c*(*s*) will be:

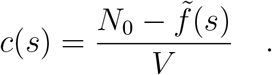

By defining 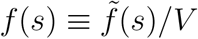, we can re-write the above equation as:

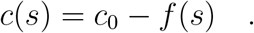

In consequence, applying equation (1), the time evolution of the reaction rate through the different division rounds of the cleavage is:

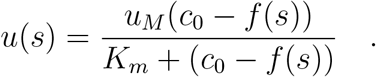

The *Cell Cycle Length* (CCL), referred in the equations as *L*, is the time spent for a division round, which is exactly the inverse than the reaction rate, i.e. *L*(*s*) = 1*/u*(*s*). Therefore, by defining the constant *L*_*µ*_ as *L*_*µ*_ = 1*/u*_*M*_, one express the evolution of the CCL *L*(*t*) as:

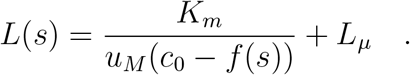

The above equation characterizes a hyperbolic growth with a singularity reachable in finite time *s*^*^. In particular, at:

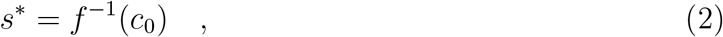

all the resources have been consumed and the CCL goes to ∞ —the cell does not duplicate anymore. We can briefly redefine terms to get a better impression of the dynamics. By rescaling the time variable as:

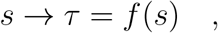

one can express *c*_0_ = *f* (*s*^*^) ≡ *τ*_*G*_ and, therefore, the evolution of *L*(*τ*) as:

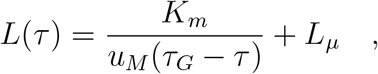

Which, by defining Λ = *K*_*m*_*/*(*u*_*M*_) is the solution of the initial value problem:

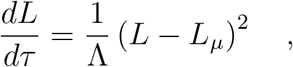

with initial conditions *L*(0) = Λ*/τ*_*G*_ + *L*_*µ*_. We remark that the above equation is valid being *τ* = *f* (*t*) *any* monotonously increasing, positive function *f* ∈ *C*^1^ such that *f* (0) = 0 and lim_*t*→∞_ *f* (*t*) = ∞. In consequence, *it is expected that the system will run into a singular behaviour in finite time*. We nevertheless, according to the data, make the assumption that the consumption of resources is such that *τ* ≈ *βs*. This will only rescale the constant Λ, being now defined as Λ = *K*_*m*_*/*(*βu*_*M*_), leading to:

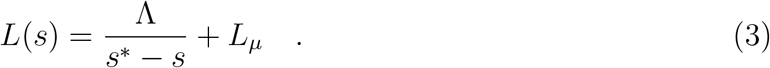

### B. Regularized duplication speed

The above equations describe an scenario where a priori there can be an explosion of the solution, since the hyperbolic nature of the described growth runs towards a singularity in finite time. Biologically, this singularity actually implies stopping the division rounds and, therefore, the death of the developing organism. To be able to safely manipulate the observed quantities, it is necessary to define a variable that avoids such singular behaviours and then, perform mapping towards the actual observables in a controlled way. To that end, we define the *reduced* rate, 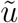, as:

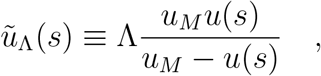

leading to:

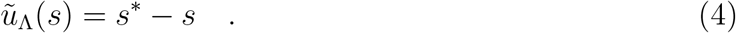

Thus, we can explicitly write *L* in terms of 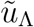 as:

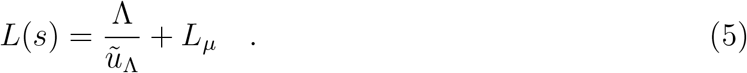

This relation will be useful when deriving the equations for the stochastic version of the problem.

### C. Relation Volume/Cell Cycle Length

Now we want to find a relation between *L* and the volume of the cells obtained by successive partitions of the cleavage. The volume of the cell at the *s*-th generation will be:

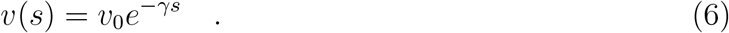

If we assume total volume conservation, 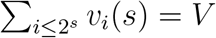, then *γ* = log 2. From the equation of the volume, one can infer the division round *s* as:

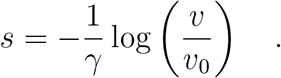

If we plug this into the equation governing the evolution of *L*, i.e., equation (3), we obtain:

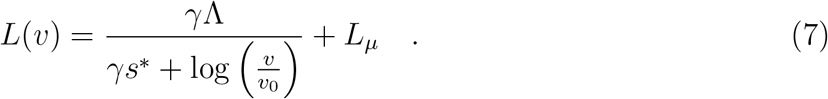

Therefore, we proved that the trend of the relation of the CCL versus cell volume is: *L*(*v*) ∝ (log *v*)^−1^. In consequence, the reduced rate 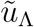, will depend on the volume, according to equation (4) as:

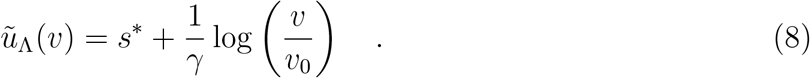

*Note: for large ranges of v, the function* (log *v*)^−1^ *may resemble v*^−*α*^, *with α* ≈ 1*/*4.

## II. STOCHASTIC DYNAMICS

We now consider that the division rounds are performed in a non-totally homogeneous way. This leads to a stochastic scenario, in which the volume of a cell *V* is not divided in totally two exactly equal volumes *V/*2, *V/*2 but to:

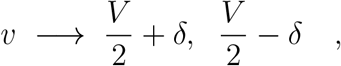

in which *δ* is a random fluctuation assumed to be small. Here we make the assumption that the volume is proportional to the available of net resources.

### A. Evolution of the cell volume

If we now follow a cell in successive generations, we can consider volume as a random variable *V* following a stochastic differential equation analogous to the one for the geometric Brownian motion with negative drift:

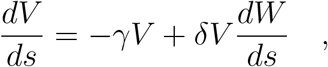

where *dW/ds* is the standard Wiener differential and *δ* a (small) noise amplitude. Recall that, in the case of perfect reductive divisions, *γ* = log 2, as above. The above stochastic differential equation has as solution the following random variable:

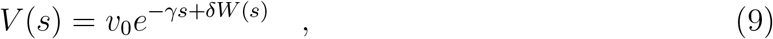

where *W* (*s*) is a Brownian random variable with mean 0 and variance 1 and, where, in addition, we made the following approximation, for the sake of simplicity: Under the assumption that *δ* ≪ 1:

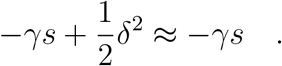

### B. Equation of evolution for *L*

We now want to account for the burst in the de-synchronization observed close to *s*^*^. We take the evolution of volume as described in equation (9) and we plug it in into equation (8) and re-write 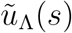 as a random process:

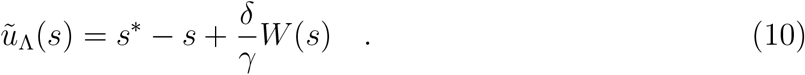

Interestingly, the above random variable describes a standard Brownian motion with linear drift, being the solution of the following SDE:

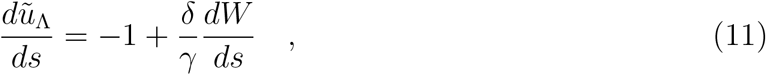

with initial value *s*(0) = *s*^*^ with probability 1. For this equation, 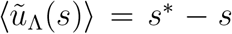 and 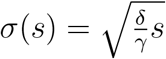.

To study the evolution of the (now stochastic variable) *L*, we make use of the relation between *L* and *u* found in equation (5) and we will consider *L* function of 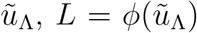. We now apply the derivation schema based on Ito’s lemma [2]:

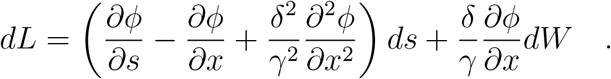

According to equation (5), *ϕ*(*x*) = Λ*/x* + *L*_*µ*_, so:

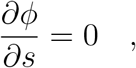

The remaining terms will lead to the following sde for *L*:

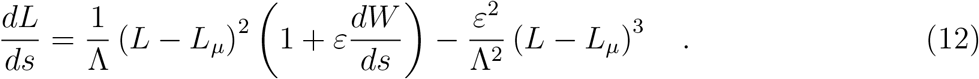

in which now *ε* = *δ/γ*. Recall that the above derivation is purely formal, because this equation leads to singular points. In order to avoid the blow-up of the solution, we introduce a *maximum CCL L*_*M*_, to avoid the divergence to ∞ and grasp the empirical fact that there is an upper bound on CCL —otherwise, the system would die. We therefore define a threshold function like:

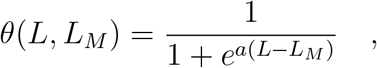

and rewrite equation (12) as:

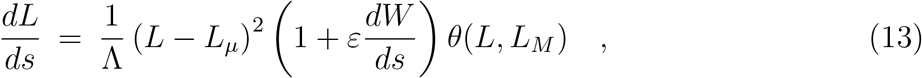

where we neglected the term:

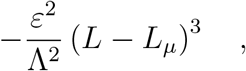

because, when limiting the growth, we assume that this high order term will not have a relevant contribution. Numerical simulations show that this is indeed the case. In addition, in order to reduce the amount of free parameters, we observe that one can define 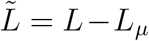, and the equation is simplified to:

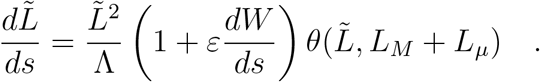

Since *L*_*M*_ is inferred from he data, one can redefine the constant as 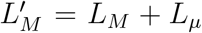. Numerical simulations show that, in the range we move in, with the proper choice of boundary conditions one can integrate equation (13) as:

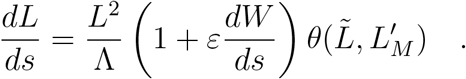

## III. CENTER-BASED FOAM MODEL

In this section we describe the equation of motion of a single-cell, modeled as a spherical particle defined by a point ***x*** and a radius *R*. In this approach, cells are represented as foam bubbles with a surface tension *γ*_*cm*_ that represents the cell’s actomyosin cortex tension, and adhesive tension *ω* that includes the reduction of actomyosin tension at a cell-cell interface *γ*_*cc*_ = *γ*_*cm*_ − *ω* (Fig. 1A) [3]. (Or equivalently the alpha-value defined as *α* = 1 − *ω/γ*_*cm*_). Using this approach, the contact angle between two cells can be computed from the tension balance at the contact point as

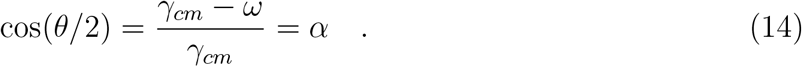

Assuming an overdamped environment, where viscous forces dominate over inertial forces, we can neglect inertial forces [4].

### A. Cell-cell contact force

The free energy of cell *i* in contact with a neighboring cell *j* can be written as

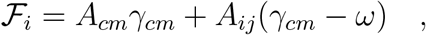

where *A*_*cm*_ is the free area of the cell in contact with the medium, and *A*_*ij*_ is the contact area with the neighboring cell. Assuming the cell volume 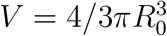 is conserved, we can calculate these areas in function of the overlap *δ* (the height of the overlapping spherical cap) between the two spheres

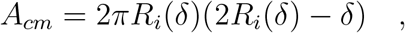

and

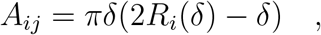

where volume conservation is represented by

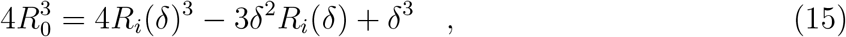

with *R*_0_ the free radius of the cell, and *R*_*i*_(*δ*) the instantaneous radius of the cell that contacts neighboring cells. So we write the free energy in function of the overlap as

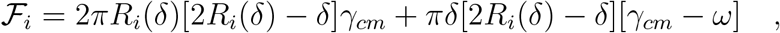

and rewrite to

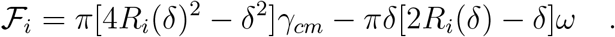

The force on cell *i* from contact with cell *j* is given by the gradient of the energy

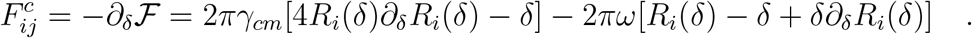

We take the derivative *∂*_*δ*_ of the volume conservation constraint, see Eq. 15, as

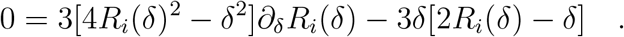

From this, we can calculate the derivative of the cell radius as

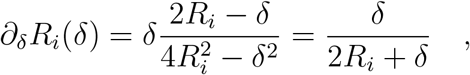

which is always positive and zero at the point of contact *δ* = 0. Finally, we can write the contact force as

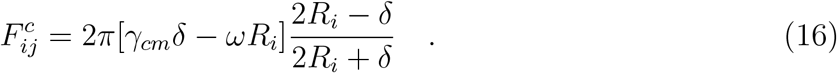

This force on cell *i* with position ***x***_*i*_ in contact with cell *j* with position ***x***_*j*_ is applied in the direction of the contact

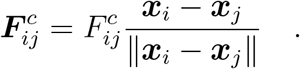

At equilibrium (Eq. 16,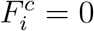), the overlap is given by *δ* = *R*_*i*_*ω/γ*_*cm*_, that can be written as a function of contact angle as cos(*θ/*2) = (*R*_*i*_ − *δ*)*/R*_*i*_ = 1 − *ω/γ* = *α*. This is exactly Eq. 14, meaning that the contact model reproduces the expected behavior of the foam contacts (Fig. 1B).

### B. Cytokinesis

Cytokinesis is modeled using a similar approach previously described by [5]. During cy-tokinesis, the mother cell is split into two daughter cells, that are represented as overlapping spheres with a small offset in the direction of division. We assume that the total volume of the two emerging daughter cells remains equal to the volume of the mother cell *V*_0_ during cytokinesis. Therefore, the radius of the daughter cells *R*_*d*_ can be computed in function of the distance *d* between the two daughter cells as

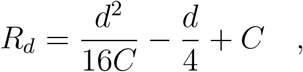

where *C* is defined as

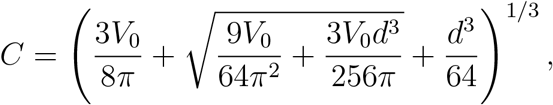

such that after division (*d* = 2*R*_*d*_) the volume of the daughter cells is half that of their mother cell 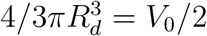. To split the daughter cells, a division force between the daughter cells is applied

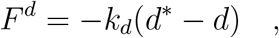

where *k*_*d*_ is the stiffness of the contact and *d*^*^ is the reference distance. The force is applied in the direction of the vector connecting the centers of the two cells. The reference distance increases linearly in time during cytokinesis

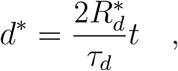

with 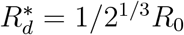 the final radius of the daughter cell, *τ*_*d*_ the duration of cytokinesis and *t* the time since the onset of cytokinesis.

**FIG. 1:**
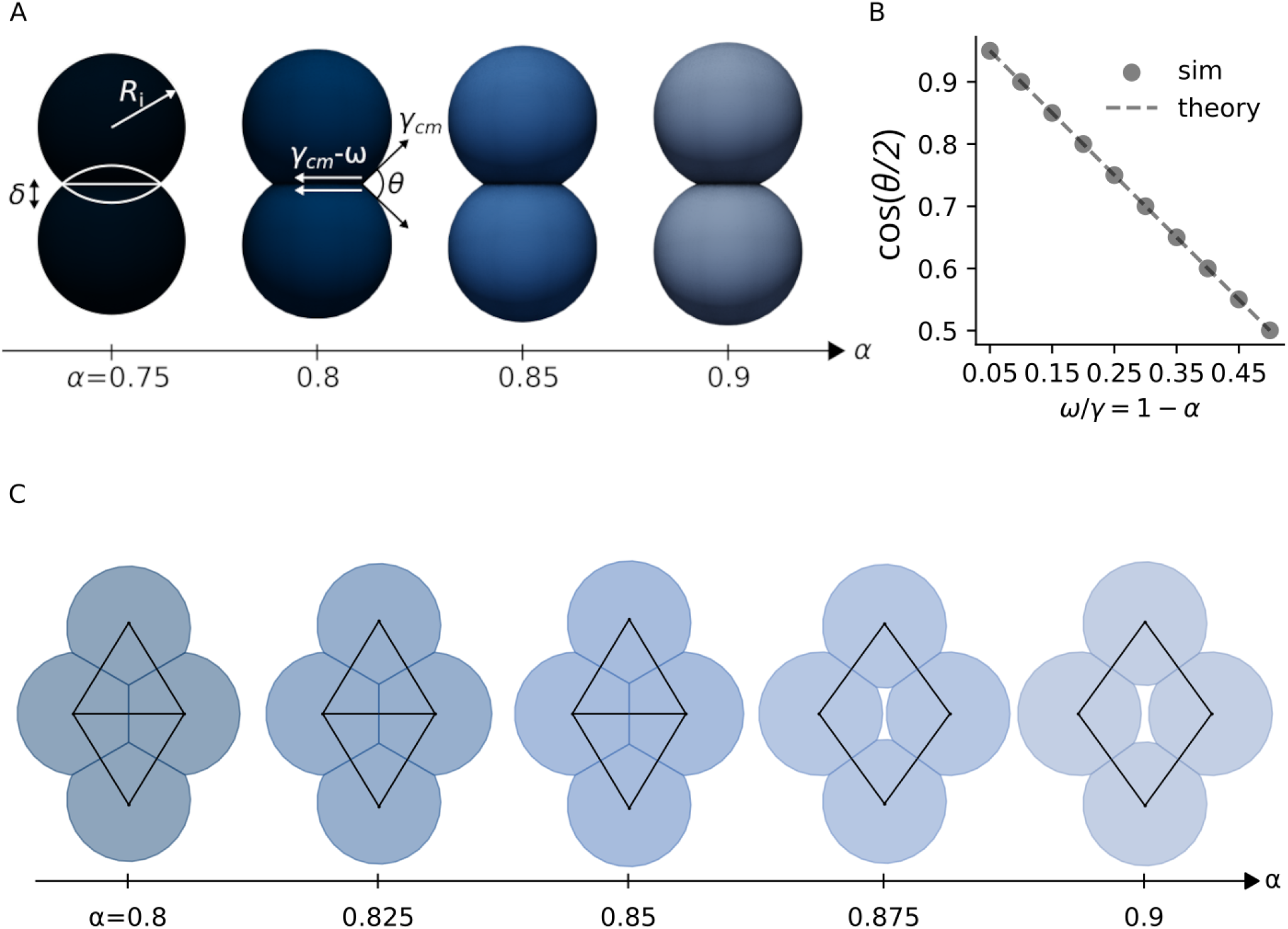
Contact model of the foam-like cells. (A) Example of cell-cell contact between two foam-like cells with surface tension *γ*_*cm*_ and adhesive tension *ω*, where *R*_*i*_ is the radius of the cell, and *δ* is the spherical overlap between the two contacting cells. (B) The contact angle between two cells in function of *α* = 1 − *ω/γ* compared to theory cos(*θ/*2) = *α*. (C) Network between 4 cells that changes from rigid to floppy in function of *α* predicted by the foam model.

**FIG. 2:**
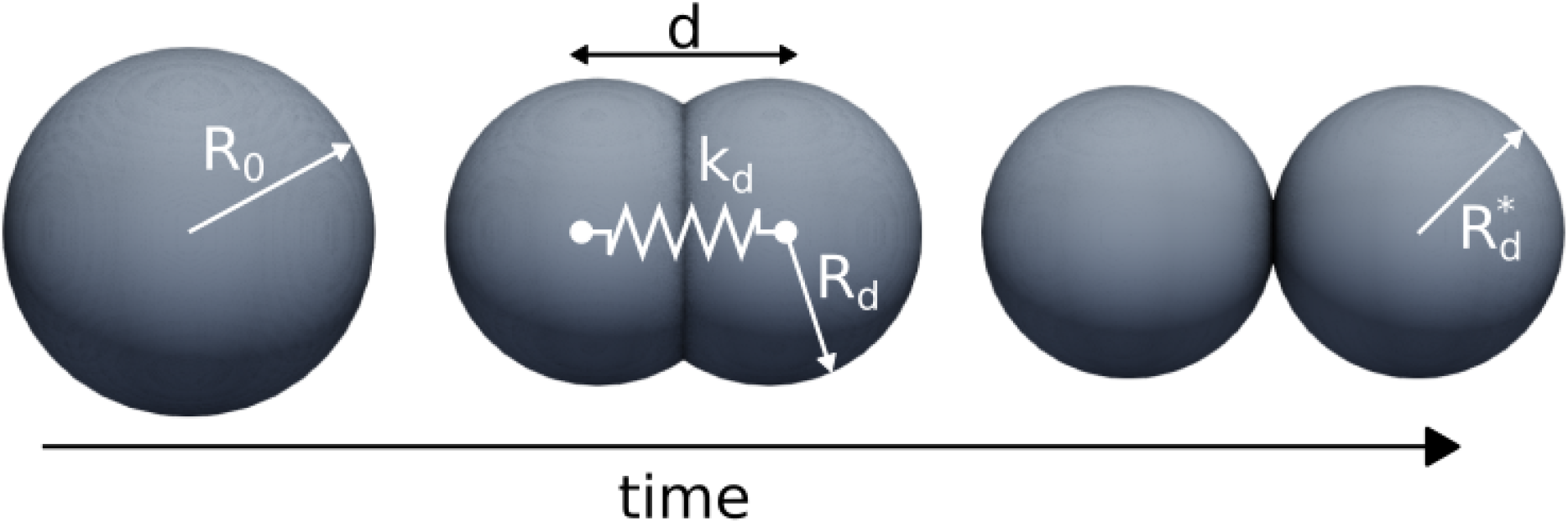
Contact model for dividing cell with constant volume 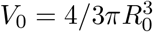. A spring with stiffness *k*_*d*_ is added between the two daughter cells. The resting length of the spring is then slowly increased, untill the daughter cells are completely separated.

### C. Damping forces

Medium damping and cell-cell friction are introduced primarily as numerical relaxation parameters rather than as direct representations of physical material properties. Their role is to provide gradual mechanical relaxation toward equilibrium, where cell-cell friction, unlike medium damping, avoids introducing artificial lag with respect to convective motion of the tissue. In practice, these damping coefficients are chosen such that the associated mechanical relaxation timescale remains sufficiently short compared to the cell cycle timescale (see Table I), ensuring quasi-static mechanical behavior. Friction between cells with relative velocity ***v***_*i*_ − ***v***_*j*_ is modeled as a viscous damping force

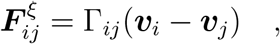

with friction tensor

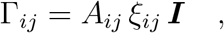

where *ξ*_*ij*_ represents the friction coefficient associated with the contact between cells *i* and *j*, and *A*_*ij*_ is the contact area. Finally, medium damping is given by

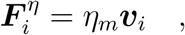

with *η*_*m*_ the effective damping coefficient of the surrounding medium.

**TABLE I:** Model parameters. Contact parameters *R*_*ij*_ and *ξ*_*ij*_ are computed for all contacts between cell *i* and cell *j*. All dissipative parameters (*ξ*_*ij*_ and *η*_*m*_ are chosen sufficiently low so the system behaves quasi-static at the timescale of the cell cycle (see main paper).

| Parameter | Symbol | Value | Unit |
| --- | --- | --- | --- |
| Initial cell radius | $R_0$ | 280 | $\mu\text{m}$ |
| Cell-medium tension | $\gamma_{cm}$ | 1 | $\text{nN } \mu\text{m}^{-1}$ |
| Adhesive tension | $\omega$ | $0.08 - 0.2$ | $\text{nN } \mu\text{m}^{-1}$ |
| Division stiffness | $k_d$ | 20 | $\text{nN } \mu\text{m}^{-1}$ |
| Average radius | $R_{ij}$ | $(R_i + R_j)/2$ | $\mu\text{m}$ |
| Cell-cell friction | $\xi_{ij}$ | $0.2k_d\Delta t/R_{ij}^2$ | $\text{kPa s } \mu\text{m}^{-1}$ |
| Medium viscosity | $\eta_m$ | 20 | $\text{nN s } \mu\text{m}^{-1}$ |
| Timestep | $\Delta t$ | 0.5 | s |

### D. Equation of motion

For each cell *i* in contact with cells *j* and daughter cell *k*, we write the equation of motion

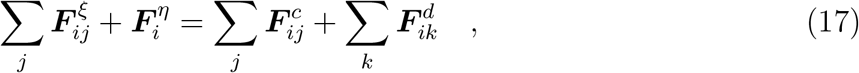

where the left-hand side denotes all friction forces that linearly depend on velocities ***v*** of the cells. For all *N* particles, we rewrite Eq. 17 as a system of equations

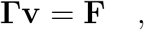

with **Γ** a 3*N* × 3*N* sparse definite matrix containing all friction components, **v** a 3*N* × 1 matrix with the velocities of all cells, and **F** a 3*N* × 1 matrix containing contact and division forces. At each timestep, we solve the system of equations for **v** using the conjugate gradient method. The positions of the cells are updated using a semi-implicit Euler scheme

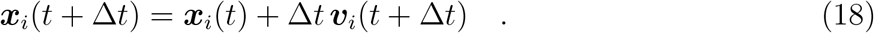

### E. Model parameters

## References

1 Mao, Y. & Wickström, S. A. Mechanical state transitions in the regulation of tissue form and function. Nat Rev Mol Cell Biol 25, 654–670, doi:10.1038/s41580-024-00719-x (2024).

2 Corominas-Murtra, B. & Petridou, N. I. Viscoelastic Networks: Forming Cells and Tissues. Frontiers in Physics 9, doi:10.3389/fphy.2021.666916 (2021).

3 Petridou, N. I. & Heisenberg, C. P. Tissue rheology in embryonic organization. EMBO J 38, e102497, doi:10.15252/embj.2019102497 (2019).

4 Rustarazo-Calvo, L., Soans, K. G. & Petridou, N. I. Tissue phase transitions in development: more than just mechanics. Development 153, doi:10.1242/dev.205219 (2026).

5 Bi, D., Lopez, J. H., Schwarz, J. M. & Manning, M. L. A density-independent rigidity transition in biological tissues. Nature Physics 11, 1074–1079, doi:10.1038/nphys3471 (2015).

6 Petridou, N. I., Corominas-Murtra, B.Heisenberg, C.-P. & Hannezo, E. Rigidity percolation uncovers a structural basis for embryonic tissue phase transitions. Cell 184, 1914–1928.e1919, doi:10.1016/j.cell.2021.02.017 (2021).

7 Kim, S., Pochitaloff, M., Stooke-Vaughan, G. A. & Campàs, O. Embryonic tissues as active foams. Nature Physics, doi:10.1038/s41567-021-01215-1 (2021).

8 Rustarazo-Calvo, L. et al. Adhesion-driven rigidity transition decoupled from density-driven jamming triggers epithelial organization in embryonic tissues. Nature Physics, doi:10.1038/s41567-026-03276-6 (2026).

9 Cugliandolo, L. F. The effective temperature. Journal of Physics A: Mathematical and Theoretical 44, 483001, doi:10.1088/1751-8113/44/48/483001 (2011).

10 Snijder, B. & Pelkmans, L. Origins of regulated cell-to-cell variability. Nature Reviews Molecular Cell Biology 12, 119–125, doi:10.1038/nrm3044 (2011).

11 Priya, R. et al. Tension heterogeneity directs form and fate to pattern the myocardial wall. Nature 588, 130–134, doi:10.1038/s41586-020-2946-9 (2020).

12 Fabrèges, D. et al. Temporal variability and cell mechanics control robustness in mammalian embryogenesis. Science 386, eadh1145, doi:10.1126/science.adh1145 (2024).

13 Ohnishi, Y. et al. Cell-to-cell expression variability followed by signal reinforcement progressively segregates early mouse lineages. Nat. Cell Biol. 16, 27–37, doi:10.1038/ncb2881 (2014).

14 Serra, D. et al. Self-organization and symmetry breaking in intestinal organoid development. Nature 569, 66–72, doi:10.1038/s41586-019-1146-y (2019).

15 Balázsi, G., van Oudenaarden, A. & Collins, J. J. Cellular decision making and biological noise: from microbes to mammals. Cell 144, 910–925, doi:10.1016/j.cell.2011.01.030 (2011).

16 Novák, B. & Tyson, J. J. Design principles of biochemical oscillators. Nat Rev Mol Cell Biol 9, 981–991, doi:10.1038/nrm2530 (2008).

17 Metallo, C. M. & Vander Heiden, M. G. Understanding metabolic regulation and its influence on cell physiology. Mol Cell 49, 388–398, doi:10.1016/j.molcel.2013.01.018 (2013).

18 Balaban, N. Q., Merrin, J., Chait, R., Kowalik, L. & Leibler, S. Bacterial Persistence as a Phenotypic Switch. Science 305, 1622–1625, doi:10.1126/science.1099390 (2004).

19 Mura, M., Feillet, C., Bertolusso, R., Delaunay, F. & Kimmel, M. Mathematical modelling reveals unexpected inheritance and variability patterns of cell cycle parameters in mammalian cells. PLoS Comput. Biol. 15, e1007054, doi:10.1371/journal.pcbi.1007054 (2019).

20 Sandler, O. et al. Lineage correlations of single cell division time as a probe of cell-cycle dynamics. Nature 519, 468–471, doi:10.1038/nature14318 (2015).

21 Godard, B. G. & Heisenberg, C.-P. Cell division and tissue mechanics. Curr Opin Cell Biol 60, 114–120, 10.1016/j.ceb.2019.05.007 (2019).

22 Schindler-Johnson, M. & Petridou, N. I. Collective effects of cell cleavage dynamics. Front. Cell Dev. Biol. 12, 1358971, doi:10.3389/fcell.2024.1358971 (2024).

23 Petridou, N. I., Grigolon, S., Salbreux, G., Hannezo, E. & Heisenberg, C. P. Fluidization-mediated tissue spreading by mitotic cell rounding and non-canonical Wnt signalling. Nat Cell Biol 21, 169–178, doi:10.1038/s41556-018-0247-4 (2019).

24 Bocanegra-Moreno, L., Singh, A., Hannezo, E., Zagorski, M. & Kicheva, A. Cell cycle dynamics control fluidity of the developing mouse neuroepithelium. Nat. Phys., doi:10.1038/s41567-023-01977-w (2023).

25 Ranft, J. et al. Fluidization of tissues by cell division and apoptosis. Proc. Natl. Acad. Sci. U. S. A. 107, 20863–20868, doi:10.1073/pnas.1011086107 (2010).

26 Firmino, J., Rocancourt, D., Saadaoui, M., Moreau, C. & Gros, J. Cell Division Drives Epithelial Cell Rearrangements during Gastrulation in Chick. Dev. Cell 36, 249–261, doi:10.1016/j.devcel.2016.01.007 (2016).

27 Saadaoui, M., Rocancourt, D., Roussel, J., Corson, F. & Gros, J. A tensile ring drives tissue flows to shape the gastrulating amniote embryo. Science 367, 453–458, doi:10.1126/science.aaw1965 (2020).

28 Garcia, S. et al. Physics of active jamming during collective cellular motion in a monolayer. Proc Natl Acad Sci U S A 112, 15314–15319, doi:10.1073/pnas.1510973112 (2015).

29 Miroshnikova, Y. A. et al. Adhesion forces and cortical tension couple cell proliferation and differentiation to drive epidermal stratification. Nat. Cell Biol. 20, 69–80, doi:10.1038/s41556-017-0005-z (2018).

30 Santos-Durán, G. N., Cooper, R. L., Jahanbakhsh, E., Timin, G. & Milinkovitch, M. C. Self-organized patterning of crocodile head scales by compressive folding. Nature 637, 375–383, doi:10.1038/s41586-024-08268-1 (2025).

31 Pauklin, S. & Vallier, L. The cell-cycle state of stem cells determines cell fate propensity. Cell 155, 135–147, doi:10.1016/j.cell.2013.08.031 (2013).

32 Gruenheit, N. et al. Cell Cycle Heterogeneity Can Generate Robust Cell Type Proportioning. Dev. Cell 47, 494–508.e494, doi:10.1016/j.devcel.2018.09.023 (2018).

33 Campinho, P. et al. Tension-oriented cell divisions limit anisotropic tissue tension in epithelial spreading during zebrafish epiboly. Nat Cell Biol 15, 1405–1414, doi:10.1038/ncb2869 (2013).

34 Sallé, J. & Minc, N. Cell division geometries as central organizers of early embryo development. Seminars in Cell & Developmental Biology 130, 3–11, 10.1016/j.semcdb.2021.08.004 (2022).

35 Edwards, S. F. & Oakeshott, R. B. S. Theory of powders. Physica A 157, 1080–1090, doi:10.1016/0378-4371(89)90034-4 (1989).

36 Shishvan, S. S., Vigliotti, A. & Deshpande, V. S. The homeostatic ensemble for cells. Biomech Model Mechanobiol 17, 1631–1662, doi:10.1007/s10237-018-1048-1 (2018).

37 Deshpande, V., DeSimone, A., McMeeking, R. & Recho, P. Chemo-mechanical model of a cell as a stochastic active gel. J. Mech. Phys. Solids 151, 104381, doi:10.1016/j.jmps.2021.104381 (2021).

38 Eldar, A. & Elowitz, M. B. Functional roles for noise in genetic circuits. Nature 467, 167–173, doi:10.1038/nature09326 (2010).

39 Hallatschek, O. et al. Proliferating active matter. Nat Rev Phys, 1–13, doi:10.1038/s42254-023-00593-0 (2023).

40 Salazar-Roa, M. & Malumbres, M. Fueling the cell division cycle. Trends Cell Biol. 27, 69–81, doi:10.1016/j.tcb.2016.08.009 (2017).

41 Ferrell, James E., Tsai Tony Y.-C. & Yang, Q. Modeling the Cell Cycle: Why Do Certain Circuits Oscillate? Cell 144, 874–885, 10.1016/j.cell.2011.03.006 (2011).

42 Kane, D. A., Warga, R. M. & Kimmel, C. B. Mitotic domains in the early embryo of the zebrafish. Nature 360, 735–737, doi:10.1038/360735a0 (1992).

43 Mohammadi, F. et al. A lineage tree-based hidden Markov model quantifies cellular heterogeneity and plasticity. Commun. Biol. 5, 1258, doi:10.1038/s42003-022-04208-9 (2022).

44 Adrián Aguirre-Tamaral, J. R., Magdalena Schindler-Johnson, Jun-Ru Lee, Daniel R. Amor, Nicoletta I. Petridou, Bernat Corominas-Murtra. A hyperbolic cell cycle law for early embryonic developmental timing. arXiv, doi: 10.48550/arXiv.2605.13234 (2026).

45 Kane, D. A. & Kimmel, C. B. The zebrafish midblastula transition. Development 119, 447–456 (1993).

46 Farrell, J. A. & O’Farrell, P. H. From Egg to Gastrula: How the Cell Cycle Is Remodeled During the Drosophila Mid-Blastula Transition. Annual Review of Genetics 48, 269–294, doi:10.1146/annurev-genet-111212-133531 (2014).

47 O’Farrell, P. H., Stumpff, J. & Su, T. T. Embryonic cleavage cycles: how is a mouse like a fly? Current biology : CB 14, R35–R45, doi:10.1016/j.cub.2003.12.022 (2004).

48 Kimmel, C. B., Ballard, W. W., Kimmel, S. R., Ullmann, B. & Schilling, T. F. Stages of embryonic development of the zebrafish. Dev. Dyn. 203, 253–310, doi:10.1002/aja.1002030302 (1995).

49 Kane, D. A. in Methods in Cell Biology Vol. 59 (eds H. William Detrich, Monte Westerfield, & Leonard I. Zon) 11–26 (Academic Press, 1998).

50 Jacobs, D. J. & Thorpe, M. F. Generic rigidity percolation: The pebble game. Phys. Rev. Lett. 75, 4051–4054, doi:10.1103/PhysRevLett.75.4051 (1995).

51 Jacobs, D. J. & Thorpe, M. F. Generic rigidity percolation in two dimensions. Phys. Rev. E Stat. Phys. Plasmas Fluids Relat. Interdiscip. Topics 53, 3682–3693, doi:10.1103/physreve.53.3682 (1996).

52 Maxwell, J. C. I. —On Reciprocal Figures, Frames, and Diagrams of Forces. Transactions of the Royal Society of Edinburgh 26, 1–40, doi:10.1017/S0080456800026351 (1870).

53 Hayden, L., Hur, W., Vergassola, M. & Di Talia, S. Manipulating the nature of embryonic mitotic waves. Curr. Biol. 32, 4989–4996.e4983, doi:10.1016/j.cub.2022.10.014 (2022).

54 Deneke, V. E., Melbinger, A., Vergassola, M. & Di Talia, S. Waves of Cdk1 Activity in S Phase Synchronize the Cell Cycle in Drosophila Embryos. Dev. Cell 38, 399–412, doi:10.1016/j.devcel.2016.07.023 (2016).

55 Mishra, N., Li, Y. I., Hannezo, E. & Heisenberg, C.-P. Geometry-driven asymmetric cell divisions pattern cell cycles and zygotic genome activation in the zebrafish embryo. Nature Physics 22, 139–150, doi:10.1038/s41567-025-03122-1 (2026).

56 Zhang, M., Skirkanich, J., Lampson, M. A. & Klein, P. S. in Vertebrate Development: Maternal to Zygotic Control (eds Francisco Pelegri, Michael Danilchik, & Ann Sutherland) 441–487 (Springer International Publishing, 2017).

57 Collart, C., Allen, G. E., Bradshaw, C. R., Smith, J. C. & Zegerman, P. Titration of Four Replication Factors Is Essential for the Xenopus laevis Midblastula Transition. Science 341, 893–896, doi:10.1126/science.1241530 (2013).

58 Vastag, L. et al. Remodeling of the metabolome during early frog development. PLoS One 6, e16881, doi:10.1371/journal.pone.0016881 (2011).

59 Amodeo, A. A., Jukam, D., Straight, A. F. & Skotheim, J. M. Histone titration against the genome sets the DNA-to-cytoplasm threshold for the Xenopus midblastula transition. Proceedings of the National Academy of Sciences 112, E1086–E1095, doi:10.1073/pnas.1413990112 (2015).

60 Liu, B., Winkler, F., Herde, M.Witte, C.-P. & Großhans, J. A Link between Deoxyribonucleotide Metabolites and Embryonic Cell-Cycle Control. Curr. Biol. 29, 1187–1192.e1183, doi:10.1016/j.cub.2019.02.021 (2019).

61 Chari, S., Wilky, H., Govindan, J. & Amodeo, A. A. Histone concentration regulates the cell cycle and transcription in early development. Development 146, doi:10.1242/dev.177402 (2019).

62 Shindo, Y. & Amodeo, A. A. Dynamics of Free and Chromatin-Bound Histone H3 during Early Embryogenesis. Curr. Biol. 29, 359–366.e354, doi:10.1016/j.cub.2018.12.020 (2019).

63 Dutta, A. & Sinha, D. K. Zebrafish lipid droplets regulate embryonic ATP homeostasis to power early development. Open Biol. 7, doi:10.1098/rsob.170063 (2017).

64 Kilwein, M. D., Dao, T. K. & Welte, M. A. Drosophila embryos allocate lipid droplets to specific lineages to ensure punctual development and redox homeostasis. PLoS Genet. 19, e1010875, doi:10.1371/journal.pgen.1010875 (2023).

65 Farrell, J. A., Shermoen, A. W., Yuan, K. & O’Farrell, P. H. Embryonic onset of late replication requires Cdc25 down-regulation. Genes Dev. 26, 714–725, doi:10.1101/gad.186429.111 (2012).

66 Farrell, J. A. & O’Farrell, P. H. Mechanism and regulation of Cdc25/Twine protein destruction in embryonic cell-cycle remodeling. Curr. Biol. 23, 118–126, doi:10.1016/j.cub.2012.11.036 (2013).

67 Di Talia, S. et al. Posttranslational Control of Cdc25 Degradation Terminates Drosophila’s Early Cell-Cycle Program. Curr. Biol. 23, 127–132, doi:10.1016/j.cub.2012.11.029 (2013).

68 Momen-Roknabadi, A., Di Talia, S. & Wieschaus, E. Transcriptional Timers Regulating Mitosis in Early Drosophila Embryos. Cell Rep. 16, 2793–2801, doi:10.1016/j.celrep.2016.08.034 (2016).

69 Balachandra, S., Sarkar, S. & Amodeo, A. A. The Nuclear-to-Cytoplasmic Ratio: Coupling DNA Content to Cell Size, Cell Cycle, and Biosynthetic Capacity. Annu. Rev. Genet. 56, 165–185, doi:10.1146/annurev-genet-080320-030537 (2022).

70 Newport, J. & Kirschner, M. A major developmental transition in early xenopus embryos: I. characterization and timing of cellular changes at the midblastula stage. Cell 30, 675–686, doi:10.1016/0092-8674(82)90272-0 (1982).

71 Kobayakawa, Y. & Kubota, H. Y. Temporal pattern of cleavage and the onset of gastrulation in amphibian embryos developed from eggs with the reduced cytoplasm. Development 62, 83–94, doi:10.1242/dev.62.1.83 (1981).

72 Edgar, B. A., Kiehle, C. P. & Schubiger, G. Cell cycle control by the nucleo-cytoplasmic ratio in early Drosophila development. Cell 44, 365–372, doi:10.1016/0092-8674(86)90771-3 (1986).

73 Donoughe, S., Hoffmann, J., Nakamura, T., Rycroft, C. H. & Extavour, C. G. Nuclear speed and cycle length co-vary with local density during syncytial blastoderm formation in a cricket. Nat. Commun. 13, 3889, doi:10.1038/s41467-022-31212-8 (2022).

74 Arata, Y., Takagi, H., Sako, Y. & Sawa, H. Power law relationship between cell cycle duration and cell volume in the early embryonic development of Caenorhabditis elegans. Front. Physiol. 5, 529, doi:10.3389/fphys.2014.00529 (2014).

75 Arata, Y. & Takagi, H. Quantitative Studies for Cell-Division Cycle Control. Front. Physiol. 10, 1022, doi:10.3389/fphys.2019.01022 (2019).

76 Piñeros, L. et al. The nuclear-cytoplasmic ratio controls the cell cycle period in compartmentalized frog egg extract. bioRxiv, doi:10.1101/2024.07.28.605512 (2024).

77 Chen, J. et al. Atypical Cadherin Dachsous1b Interacts with Ttc28 and Aurora B to Control Microtubule Dynamics in Embryonic Cleavages. Dev. Cell 45, 376–391.e375, doi:10.1016/j.devcel.2018.04.009 (2018).

78 Knopp, K. in Theory of Functions Parts I and II, Two Volumes Bound as One, Part I 117-139 (1996).

79 Van Kampen, N. G. Stochastic processes in physics and chemistry. 3 edn, (Elsevier Science & Technology, 2007).

80 Michaelis, L., Menten, M. L., Johnson, K. A. & Goody, R. S. The original Michaelis constant: translation of the 1913 Michaelis-Menten paper. Biochemistry 50, 8264–8269, doi:10.1021/bi201284u (2011).

81 Kõivomägi, M. et al. Dynamics of Cdk1 substrate specificity during the cell cycle. Mol. Cell 42, 610–623, doi:10.1016/j.molcel.2011.05.016 (2011).

82 Jukam, D., Shariati, S. A. M. & Skotheim, J. M. Zygotic Genome Activation in Vertebrates. Developmental cell 42, 316–332, doi:10.1016/j.devcel.2017.07.026 (2017).

83 Tadros, W. & Lipshitz, H. D. The maternal-to-zygotic transition: a play in two acts. Development 136, 3033–3042, doi:10.1242/dev.033183 (2009).

84 Rathbun, L. I. et al. PLK1-and PLK4-Mediated Asymmetric Mitotic Centrosome Size and Positioning in the Early Zebrafish Embryo. Curr. Biol. 30, 4519–4527.e4513, doi:10.1016/j.cub.2020.08.074 (2020).

85 Afonso, O., Dumoulin, L., Kruse, K., and Gonzalez-Gaitan, M. Cytoplasmic flow is a cell size sensor that scales anaphase. Nat Cell Bio (2025).

86 Olivier, N. et al. Cell Lineage Reconstruction of Early Zebrafish Embryos Using Label-Free Nonlinear Microscopy. Science 329, 967–971, doi:10.1126/science.1189428 (2010).

87 Hadjivasiliou, Z. et al. Basal Protrusions Mediate Spatiotemporal Patterns of Spinal Neuron Differentiation. Dev. Cell 49, 907–919.e910, doi:10.1016/j.devcel.2019.05.035 (2019).

88 Cover, T. M. & Thomas, J. A. Elements of Information Theory. 99 edn, (John Wiley & Sons, 1991).

89 Maître, J.-L. et al. Adhesion Functions in Cell Sorting by Mechanically Coupling the Cortices of Adhering Cells. Science 338, 253–256, doi:10.1126/science.1225399 (2012).

90 Maître, J. L., Niwayama, R., Turlier, H., Nédélec, F. & Hiiragi, T. Pulsatile cell-autonomous contractility drives compaction in the mouse embryo. Nat Cell Biol 17, 849–855, doi:10.1038/ncb3185 (2015).

91 Weaire, D. & Hutzler, S. The physics of foams. (Oxford University Press, 2001).

92 Das, R., Sinha, S., Li, X., Kirkpatrick, T. R. & Thirumalai, D. Free volume theory explains the unusual behavior of viscosity in a non-confluent tissue during morphogenesis. Elife 12, doi:10.7554/eLife.87966 (2024).

93 Autorino, C. & Petridou, N. I. Critical phenomena in embryonic organization. Curr. Opin. Syst. Biol. 31, 100433, doi:10.1016/j.coisb.2022.100433 (2022).

94 Kitano, H. Biological robustness. Nature Reviews Genetics 5, 826–837, doi:10.1038/nrg1471 (2004).

95 Tran, N. H. N., Nguyen, A., Rahman, T. W. & Baetica, A.-A. Fundamental Trade-Offs in the Robustness of Biological Systems with Feedback Regulation. ACS Synth Biol 14, 1099–1111, doi:10.1021/acssynbio.4c00704 (2025).

96 Moghe, P., Hannezo, E. & Hiiragi, T. Optimality as a framework for understanding developmental robustness. Trends Cell Biol. 36, 246–256 (2025).

97 Hannezo, E. & Heisenberg, C. P. Mechanochemical Feedback Loops in Development and Disease. Cell 178, 12–25, doi:10.1016/j.cell.2019.05.052 (2019).

98 Schwayer, C. et al. Cell heterogeneity and fate bistability drive tissue patterning during intestinal regeneration. bioRxiv, doi:10.1101/2025.01.14.632683 (2025).

99 Salvidge, W. et al. Linage priming and cell type proportioning depends on the interplay between stochastic and deterministic factors. bioRxiv, doi:10.1101/2025.01.08.631908 (2025).

100 Stewart, P. S. & Franklin, M. J. Physiological heterogeneity in biofilms. Nat. Rev. Microbiol. 6, 199–210, doi:10.1038/nrmicro1838 (2008).

101 Ulfig, A. & Jakob, U. Redox heterogeneity in mouse embryonic stem cells individualizes cell fate decisions. Dev. Cell 59, 2118–2133.e2118, doi:10.1016/j.devcel.2024.07.008 (2024).

102 Desai, R. V. et al. A DNA repair pathway can regulate transcriptional noise to promote cell fate transitions. Science 373, eabc6506, doi:10.1126/science.abc6506 (2021).

103 Gomer, R. H. & Firtel, R. A. Cell-autonomous determination of cell-type choice in Dictyostelium development by cell-cycle phase. Science 237, 758–762, doi:10.1126/science.3039657 (1987).

104 Raj, A. & van Oudenaarden, A. Nature, nurture, or chance: stochastic gene expression and its consequences. Cell 135, 216–226, doi:10.1016/j.cell.2008.09.050 (2008).

105 Maamar, H., Raj, A. & Dubnau, D. Noise in gene expression determines cell fate in Bacillus subtilis. Science 317, 526–529, doi:10.1126/science.1140818 (2007).

106 Chang, H. H., Hemberg, M., Barahona, M., Ingber, D. E. & Huang, S. Transcriptome-wide noise controls lineage choice in mammalian progenitor cells. Nature 453, 544–547, doi:10.1038/nature06965 (2008).

107 Lord, N. D. et al. Stochastic antagonism between two proteins governs a bacterial cell fate switch. Science 366, 116–120, doi:10.1126/science.aaw4506 (2019).

108 Yanagida, A. et al. Cell surface fluctuations regulate early embryonic lineage sorting. Cell 185, 777–793.e720, doi:10.1016/j.cell.2022.01.022 (2022).

109 Johnson, S. L. & Bennett, P. Growth control in the ontogenetic and regenerating zebrafish fin. Methods Cell Biol. 59, 301–311, doi:10.1016/s0091-679x(08)61831-2 (1999).

110 Stegmaier, J. et al. Real-Time Three-Dimensional Cell Segmentation in Large-Scale Microscopy Data of Developing Embryos. Dev. Cell 36, 225–240, doi:10.1016/j.devcel.2015.12.028 (2016).

111 Schindelin, J. et al. Fiji: an open-source platform for biological-image analysis. Nat. Methods 9, 676–682, doi:10.1038/nmeth.2019 (2012).

112 Ershov, D. et al. TrackMate 7: integrating state-of-the-art segmentation algorithms into tracking pipelines. Nat. Methods 19, 829–832, doi:10.1038/s41592-022-01507-1 (2022).

113 Tinevez, J.-Y. et al. TrackMate: An open and extensible platform for single-particle tracking. Methods 115, 80–90, doi:10.1016/j.ymeth.2016.09.016 (2017).

114 Wickham, H. Ggplot2: Elegant graphics for data analysis. 2 edn, (Springer International Publishing, 2016).

115 Wickham, H. et al. Welcome to the tidyverse. Journal of Open Source Software version 1.2.0 4, 1686, doi:10.21105/joss.01686 (2019).

116 Wickham, H., François, R., Henry, L., Müller, K. & Vaughan, D. dplyr: A Grammar of Data Manipulation. version 1.0.10 (2022).

117 Machado, S., Mercier, V. & Chiaruttini, N. LimeSeg: a coarse-grained lipid membrane simulation for 3D image segmentation. BMC Bioinformatics 20, 2, doi:10.1186/s12859-018-2471-0 (2019).

118 Autorino, C. et al. Tissue rigidity phase transition shapes morphogen gradients. Nature Cell Biology, doi:10.1038/s41556-026-01954-4 (2026).

119 Stringer, C., Wang, T., Michaelos, M. & Pachitariu, M. Cellpose: a generalist algorithm for cellular segmentation. Nat. Methods 18, 100–106, doi:10.1038/s41592-020-01018-x (2021).

120 Harris, C. R. et al. Array programming with NumPy. Nature 585, 357–362, doi:10.1038/s41586-020-2649-2 (2020).

121 Kassambara, A. ggpubr: ‘ggplot2’ Based Publication Ready Plots. version 0.5.0 (2022).

## References

[1] A. Aguirre-Tamaral, J. Royer, M. Schindler-Johnson, J.-R. Lee, D. R. Amor, N. I. Petridou, and B. Corominas-Murtra, arXiv preprint arXiv:2605.13234 (2026), 2605.13234.

[2] W. Horsthemke and R. Lefever, Noise-Induced Transitions: Theory and Applications in Physics, Chemistry, and Biology, Springer Series in Synergetics (Springer Berlin Heidelberg, 2006).

[3] J.-L. Maître, H. Berthoumieux, S. F. G. Krens, G. Salbreux, F. Jülicher, E. Paluch, and C.-P. Heisenberg, Science 338, 253 (2012).

[4] E. M. Purcell, American Journal of Physics 45, 3 (1977).

[5] S. Ongenae, M. Cuvelier, J. Vangheel, H. Ramon, and B. Smeets, Frontiers in Physics 9 (2021).

